# Experiences are encoded by brainwide reprogramming of synaptome architecture

**DOI:** 10.64898/2025.12.12.693919

**Authors:** Hanan Woods, Ülkü Günar, Sarah Catherine Gillard, Beverley Notman, Kunhao Yuan, Digin Dominic, Emily Robson, Gabor Varga, Noboru H. Komiyama, Zhen Qiu, Frank Sengpiel, Seth G.N. Grant

**Author notes:** Equal contribution (Ülkü Günar, Sarah Catherine Gillard).

## Abstract

Synaptome architecture describes the spatiotemporal distribution of highly diverse excitatory synapses throughout the brain. Whether and how this architecture is impacted by experience is key to understanding its role in learning and memory. We found that environmental enrichment and monocular visual deprivation drive large-scale, type-and subtype-specific reorganisation of excitatory synapses in more than one hundred brain regions. Each experience modifies distinct subsets of synapses, with patterns aligned with protein turnover rates and connectome architecture. These reorganisations occur during development and adulthood, revealing a conserved mechanism of synaptome plasticity across the lifespan. Our findings support a population-selection model in which experience drives adaptation by selectively modifying synapse varieties, generating a distributed trace of past experiences. Our results also point to synaptome architecture as a shared framework integrating experience, lifespan changes, sleep, genetic variation and disease.

## Introduction

The vast majority of brain synapses are excitatory, and their proteomes are highly complex^1–12^ and extraordinarily diverse^13–16^. Individual synapses differ in their protein composition^13,16^, rate of protein turnover^17^ and their nanoarchitecture^18–20^. Systematic, data-driven analysis of individual synapse proteomes has led to the concept of the synaptome – the complete population of diverse synapse types in the brain – and its spatial and temporal organization, known as the synaptome architecture^16^. Single-synapse resolution mapping of excitatory synapse types and subtypes across the mouse brain has revealed that the synaptome architecture changes continuously across the lifespan^21,22^ and in response to genetic mutations^16,22^ and that sleep is essential for its maintenance^23^. Together, these findings indicate that the synaptome and synaptome architecture are genetically programmed and flexible.

Whether experience alters the synaptome architecture remains an open question. To investigate environmental influences, we used the environmental enrichment (EE) paradigm^24,25^, comparing mice raised in complex environments with those housed in standard cages (SC). To assess the impact of specific sensory input during development, we also examined mice subjected to one week of monocular deprivation (MD) during the postnatal critical period^26,27^. Synaptome mapping technology enables single-synapse analysis of protein composition, turnover and morphology across the mouse brain by imaging endogenously tagged synaptic proteins in tissue sections^16,17,21–23^. Fluorescent^16^ or self-labelling^17^ tags report protein abundance and turnover, allowing classification of excitatory synapses into types based on protein presence and into subtypes according to abundance, turnover, morphology and size, revealing extensive molecular diversity across brain regions. In this study, mice expressing PSD95-eGFP and SAP102-mKO2 were used to classify synapses into three types—Type 1 (PSD95 only), Type 2 (SAP102 only) and Type 3 (dual-labelled)—and 37 subtypes^16^. Thirty of these subtypes can be further categorised by short (SPL), medium (MPL) or long (LPL) PSD95 protein lifetime^17,22,23^.

To our knowledge, this is the first study to investigate the impact of experience during development on synapse type and subtype populations across whole brain sections. We show that distinct experiences induce selective and spatially distributed alterations in specific excitatory synapse types and subtypes, with the resulting patterns shaped by protein turnover dynamics and the underlying connectome. These findings demonstrate that experience is encoded through modifications to defined synapse populations, revealing a crucial role for synapse diversity in core brain function.

### Developmental experience shapes synaptome architecture

As a first step toward establishing whether synaptome architecture is shaped by experience we reared mice expressing PSD95-eGFP and SAP102-mKO2 from birth to three months in enriched (EE; n = 23) or standard (SC; n = 29) housing (Fig. 1A). Using our synaptome mapping pipeline, we analysed an average of 23.9 million excitatory synaptic puncta across 100 brain regions from each mouse and mapped the location of three types and 37 subtypes of excitatory synapse. The systematic nature of our analysis contrasts with prior studies that have typically sampled fewer than 5,000 synapses per mouse in a single brain region in very few animals.

**Figure 1.**
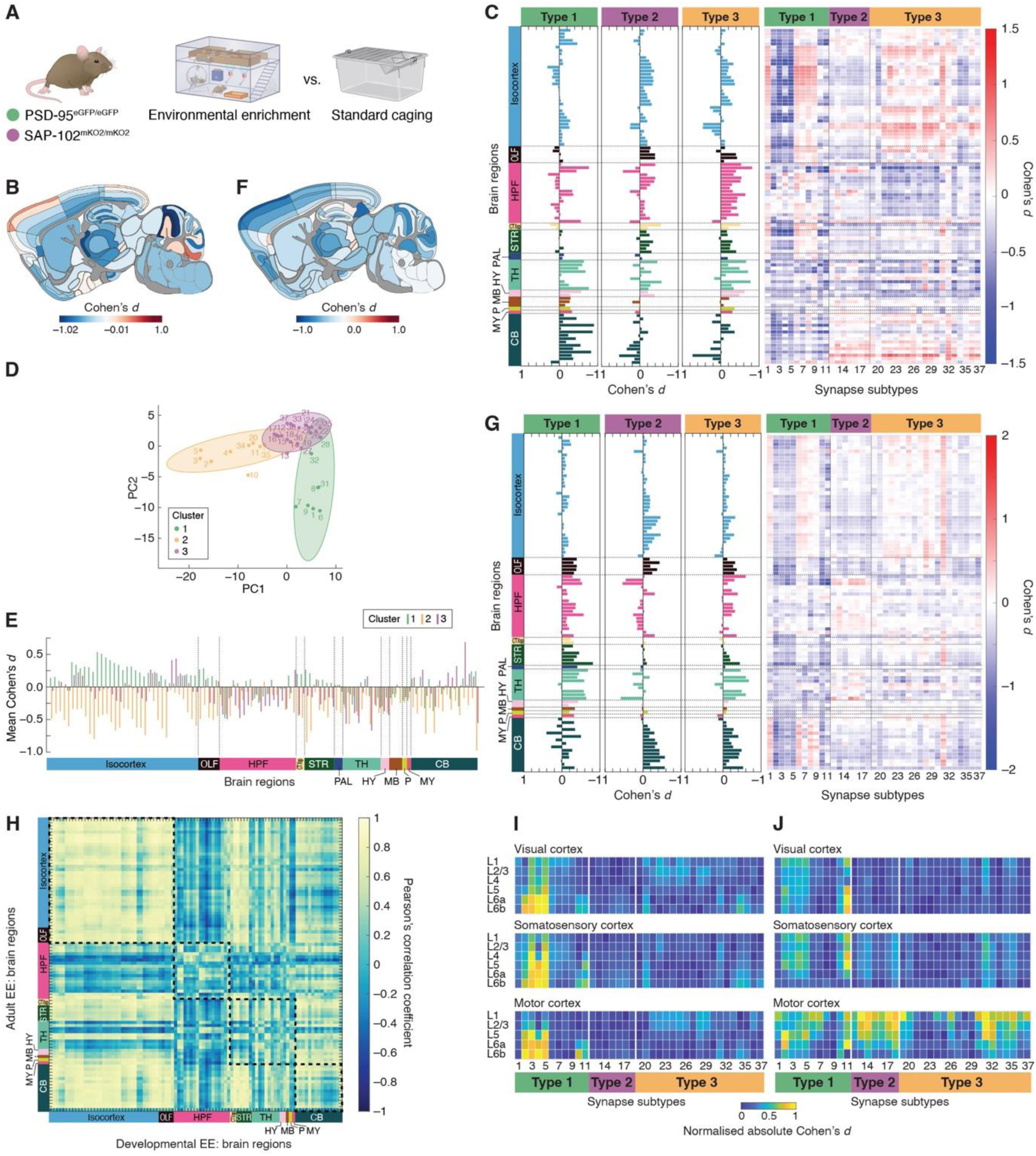
Developmental and adult environmental enrichment remodel brain synaptome architecture. **(A)** Experimental groups. *Psd95^eGFP/eGFP^*and *Sap102^mKO2/mKO2^*mice were reared under environmental enrichment (EE; *n* = 23, 12 females) or standard cage (SC; *n* = 29, 15 females) conditions for 3 months from birth. In a separate experiment, adult mice previously housed under SC conditions were transferred to EE (*n* = 20, all female) or maintained in SC (*n* = 20, all female) for 1 month. **(B, F)** Parasagittal brain maps showing Cohen’s *d* effect size values for total synapse density between EE and SC groups in juvenile EE (B) and adult EE (F) cohorts. Positive values indicate higher synapse density in EE relative to SC. **(C, G)** Regional effect sizes (Cohen’s *d*) for three synapse types (bar plots) and 37 subtypes (heatmaps), comparing EE versus SC groups in juvenile (C) and adult (G) paradigms. Effect sizes > 0 reflect enrichment-related increases. Expanded versions of these subtype heatmaps, showing complete region labels and statistical significance markers, are available in Fig. S1 and S8. **(D)** Biplot of robust principal component analysis (PCA) of subtype effect size profiles across brain regions. Subtypes are coloured by cluster identity (see key), defined via hierarchical clustering of PCA scores. **(E)** Mean Cohen’s *d* per brain subregion stratified by subtype cluster (see key). **(H)** Pearson’s correlation matrix comparing subtype-by-region Cohen’s *d* effect size matrices from juvenile and adult EE cohorts, quantifying similarity in enrichment-driven synaptome remodelling. **(I, J)** Normalized absolute Cohen’s *d* values for subtype-layer combinations in selected cortical areas (visual, somatosensory, motor cortex) for juvenile (I) and adult (J) EE paradigms. Within each cortical area, values were normalized by dividing each effect size by the maximum effect size observed across all layers and subtypes. For brain region abbreviations see supplementary material ‘Brain region list’.

Surprisingly, unlike some earlier studies^28–34^, we found EE did not lead to a greater density of excitatory synapses across the whole brain, major brain regions, or subregions (Bayesian analysis with FDR correction; Fig. 1B). Instead, a modest trend toward lower synapse density emerged across many regions, with a median decrease of 3% observed across subregions. These findings suggest that EE does not broadly promote global synaptogenesis, but instead may drive more subtle, selective remodelling within specific excitatory synapse populations.

To explore this possibility, we quantified the differences in density of synapse types and subtypes in mice raised in EE and SC (Fig. 1C). This revealed a striking differential pattern in the synaptome across the whole brain. EE resulted in a reduction in the density of Type 1 synapses in the molecular layer of the cerebellum, thalamus, and dentate gyrus (Fig. 1C). A subset of Type 1 subtypes (2–5, 10, 11) were consistently reduced in EE mice across diverse structures, including the deep cortical layers, cerebellar layers, thalamus, hypothalamus, striatum, pallidum, olfactory regions, and cortical subplate (Fig. 1C and S1). By contrast, the remaining Type 1 subtypes (1, 6-9) were increased in several regions, most notably in the cortex. Moreover, the strongest synaptic changes in the cortex were found in the deeper layers for subtypes 4 and 5 (respectively p < 0.0014 and p < 0.0061, FDR-corrected). Differences in Type 3 synapse subtype densities showed similar trends across cortical regions, including differential effects in deep and superficial layers, and between the hippocampus, thalamus, and hypothalamus, whereas Type 2 synapses were largely unaffected (Fig. 1C and S1).

To identify coordinated patterns of synaptome reprogramming in groups of subtypes that might not be evident at the individual subtype level, we applied robust principal component analysis (PCA) across all 37 synapse subtype effect size values (Fig. 1D). The first three components accounted for 43.7%, 19.3%, and 12.0% of the variance, respectively, with the stability of PC1 confirmed by bootstrap resampling (95% CI: 37.4%–67.6%) and permutation testing (p < 0.001). Hierarchical clustering of PCA scores yielded three stable clusters (Jaccard indices: 0.81, 0.68 and 0.79), which showed robust separation along the first two principal components (Kruskal-Wallis, PC1: χ² = 24.9, df = 2, p = 3.9 × 10⁻⁶; PC2: χ² = 23.4, df = 2, p = 8.1 × 10⁻⁶). Notably, Cluster 2 contained most subtypes shown above to decrease with EE (subtypes 2–5, 10, 11, 20, 34, 35; Fig. 1D) and was maximally distinct from Cluster 1 (subtypes 1, 6– 9, 28, 31, 32) along PC1. EE-dependent divergence in these clusters was most evident in the isocortex, striatum, olfactory areas, putamen, and cortical subplate (Fig. 1E). Cluster 3 subtypes largely correspond to those that showed little change in response to EE – all of the Type 2 subtypes (12-18) and the remaining Type 3 subtypes (19, 21-27, 29, 30, 33, 36, 37). Together, these findings support a model in which sensory experiences during development drive selective, type-and subtype-specific shifts in synapse populations across all brain regions.

We observed differential patterns in the synaptome architecture of male and female mice raised in the two environments (Fig. S2-S7). Although sex did not significantly interact with housing condition, and within-group comparisons between males and females revealed no major corrected differences, trends suggest that EE may exaggerate sex differences across multiple brain regions (Fig. S2-S7).

### EE remodels the synaptome architecture of the adult brain

It is generally thought that the developing brain is more plastic than the mature brain ^35,36^. To explore this issue, we asked whether exposing mature mice (6-12 months of age) for one month to the same EE used in the aforementioned experiments would induce a change in synaptome architecture. Strikingly, the same broad trends of synaptome reprogramming were seen as in the developmental EE experiment. Total synapse density was modestly reduced, and subtype-specific decreases were detected in subtypes 2–5, 10, 11, 20, 34, and 35, closely matching the EE-sensitive cluster identified in the developmental cohort (Fig.1F,G and S8). The similarity matrix shows highest similarity in the isocortex, with hippocampal, subcortical and cerebellar regions showing differential responses between developmental and adult brain (Fig. 1H).

Although there was an overall consistency of affected subtypes in both developmental and adult EE, implying that a core population of excitatory synapses remains responsive to environmental input into adulthood, there were differences within particular brain areas. For example, in the isocortex the main differential effects were in the deep layers with developmental EE, whereas it was in the superficial layers with adult EE (Fig. 1I,J). Moreover, Type 2 and 3 subtypes in the motor cortex and orbital cortex were differentially affected between developmental and adult EE paradigms (Fig. 1C,G,I,J).

### MD drives global synaptome changes

The brainwide changes in synaptome architecture induced by EE might have arisen because of input to multiple sensory pathways. We therefore explored the impact of a limited sensory deprivation, namely MD, in mice from postnatal day 25/26 to 31 (treated n = 13; control n = 14) (Fig. 2A). At the end of the deprivation period, coronal sections of both hemispheres, which included the primary and secondary visual regions among a total of 119 regions, were analysed. In each section we mapped an average of 32.5 million excitatory synaptic puncta, which contrasts with earlier studies analysing fewer than 5,000 synapses in only visual regions^37–39^.

**Figure 2.**
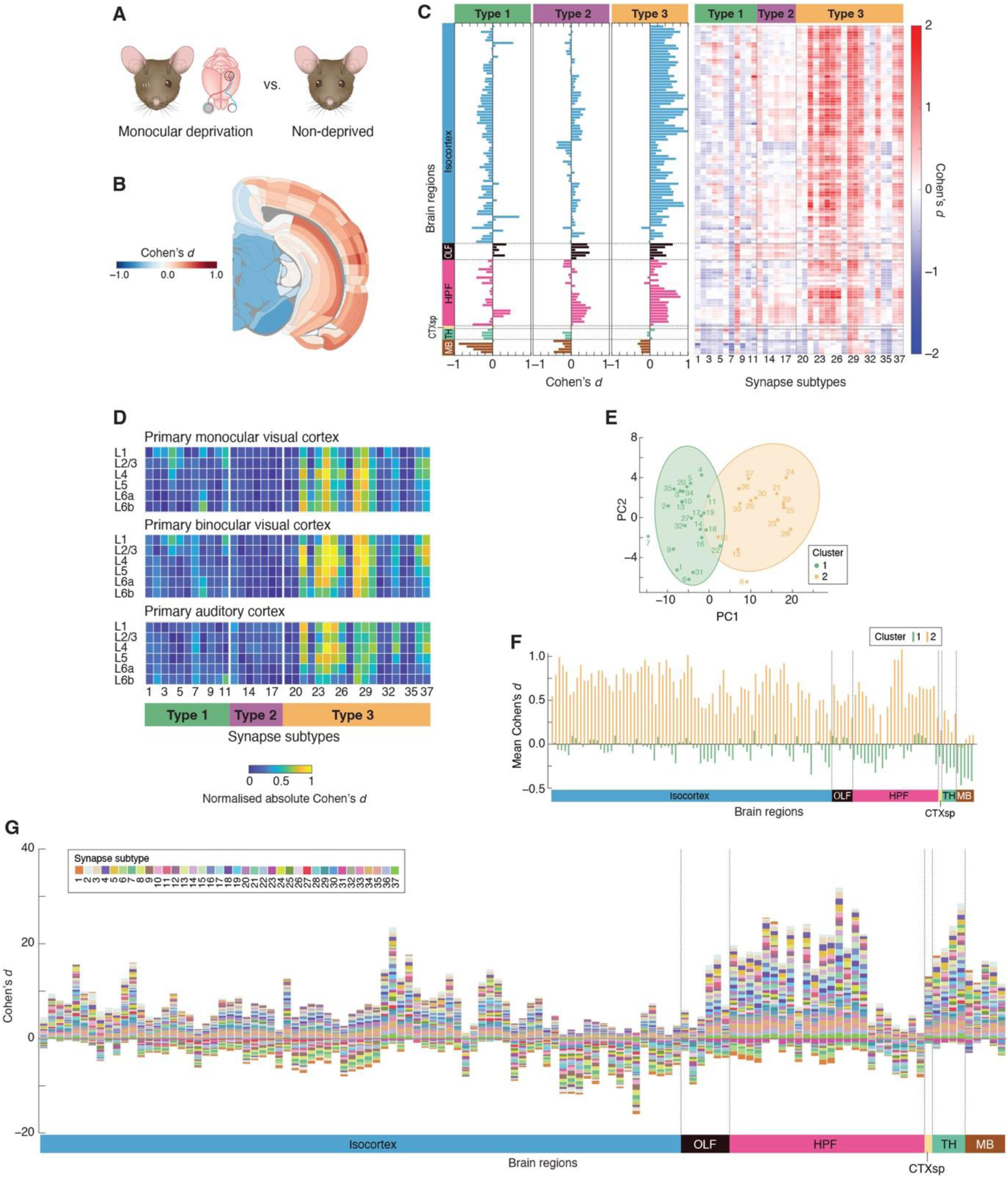
Monocular deprivation remodels synaptome architecture in visual and non-visual cortical regions. **(A)** Experimental design. Mice homozygous for *Psd95^eGFP^* and *Sap102^mKO2^* underwent monocular deprivation (MD) via right eyelid suture from postnatal day (P) 25/26 to P31 (*n* = 13, 7 female) and were compared with non-deprived controls (CTRL; *n* = 14, 7 female). The right-eye suture deprived the contralateral (left) monocular zone of the primary visual cortex of visual input and partially affected visual input to binocular visual cortex in both hemispheres. **(B)** Coronal brain map showing Cohen’s *d* effect size values for total synapse density across regions, comparing MD and CTRL groups. Positive values indicate higher synapse density in the MD group. **(C)** Regional effect sizes (Cohen’s *d*) for three synapse types (bar plots) and 37 subtypes (heatmaps), comparing MD versus CTRL. Positive values indicate MD-associated increases in synapse density. Expanded versions of the subtype heatmaps and stacked bar graph, including full region labels and statistical significance indicators, are provided in Fig. S9. **(D)** Normalized absolute Cohen’s *d* values for subtype-layer combinations in selected cortical regions: monocular and binocular zones of primary visual cortex, and primary auditory cortex. Within each region, values were normalized by dividing each effect size by the maximum observed across all layers and subtypes. **(E)** Biplot of robust PCA based on subtype effect size profiles across brain regions. Subtypes are coloured by cluster identity (see key), derived from hierarchical clustering of PCA scores. **(F)** Bar plot showing mean Cohen’s *d* per region, stratified by cluster. **(G)** Stacked bar graph of Cohen’s *d* values for 37 synapse subtypes in the contralateral (deprived) hemisphere, comparing MD and CTRL. Subregion values were normalized by dividing contralateral densities by the sum of ipsilateral and contralateral densities. Positive values indicate increased relative density in MD. For brain region abbreviations see supplementary material ‘Brain region list’.

Although one might anticipate that MD would produce synaptome changes primarily in the visual system, we observed distributed impacts broadly similar to those in response to EE. MD did not significantly alter total excitatory synapse density across the whole brain or within major brain regions (Bayesian analysis with FDR correction; Fig. 2B) but there were striking changes in the densities of particular synapse types and subtypes on a brainwide scale (Fig. 2C, S9 and S10). Type 3 synapses were consistently elevated across nearly all isocortical and hippocampal subregions in the deprived hemisphere (Fig. 2C and S9). Type 1 and 2 synapse populations exhibited smaller, region-specific changes. These effects were not confined to the visual cortex or even the deprived contralateral hemisphere. Type 3 synapse increases were mirrored in the non-deprived, ipsilateral hemisphere and extended into multiple sensory cortices, suggesting a distributed cortical response to the MD paradigm (Fig. S10). Synapse subtype analysis further revealed that specific Type 3 subtypes (21, 23–26, 28–30, 33, 36, 37) were elevated across both hemispheres in MD mice, including in auditory, somatosensory, and hippocampal regions (Fig. 2C,D, S9 and S10).

We identified coordinated patterns of synaptome reprogramming using PCA analysis across all 37synapse subtype effect size values. PC1 explained 79.6% of the variance (95% CI: 60.7–86.3%; p < 0.001), indicating a dominant and structured response to MD. Hierarchical clustering in PCA space identified two clusters (Jaccard indices: 0.524, 0.635), differing significantly in PC1 scores (Kruskal-Wallis, PC1: χ² = 25.4, df = 1, p = 4.6 × 10⁻7; PC2: χ² = 0.09, df = 1, p = 0.75). These subtype groupings differed from those identified for the EE paradigm. Here, Cluster 2 encompassed the MD-sensitive Type 3 subtypes (21, 23-26, 28-30, 33, 36, 37) as well as subtypes 8, 12 and 15, supporting their coordinated elevation across multiple regions (Fig. 2E). Whereas MD was associated with localized increases in Type 3 subtypes, particularly within isocortical and hippocampal regions, EE produced a subtle downscaling of Type 1 and 3 subtypes (Fig. 2C,F, Fig. 1C,E). Thus, EE and MD paradigms reprogram the synaptome through selective and distributed changes in distinct synapse types and subtypes.

We identified hemisphere-specific effects of MD (see methods), which selectively increased the relative density of specific synapse subtypes in the contralateral hemisphere, particularly within the monocular visual cortex, retrosplenial cortex, hippocampus, thalamus, and midbrain, and modified inter-hemisphere differences (Fig. 2G, S11-S13). As observed for EE, sex-dependent shifts in synaptome architecture were observed with MD, but these trends were not supported with significance testing (Fig. S14-S19; Bayesian analysis and FDR correction).

### Protein lifetime as a constraint on synaptome plasticity

Our findings point to a key question: why do only certain synapse subtypes undergo structural change following EE or MD? We hypothesized that the capacity for change may be governed, in part, by the rate of protein turnover in synapses. In this setting, synapse subtypes built from long-lived proteins may respond differently to experience than those composed of proteins that turn over more rapidly.

To test this, we aligned our effect size data for each of 30 Type 1 and Type 3 subtypes with PSD95 protein half-life estimates^17^. Reordering heatmaps of EE and MD effect sizes by protein lifetime (grouped into LPL, MPL and SPL) revealed striking patterns. Both EE and MD heatmaps show a vertical stripe in the LPL synapses and MD shows another stripe in both MPL and SPL synapses. These stripes show that EE preferentially downregulated LPL synapses, whereas MD was associated with increased density of MPL and SPL synapses (Fig. 3A,B and S20). This pattern was consistent across the isocortex and hippocampus for MD, but diverged in EE, where SPL synapses were suppressed in the hippocampus. In a separate study on the effect of sleep deprivation on synaptome architecture, we also observed a prominent striping pattern that separated LPL synapses ^23^. However, this pattern was the opposite of that observed for EE in that LPL synapses were upregulated after sleep deprivation (Fig. S22). Similarly, LPL synapses were upregulated in the deprived hemisphere relative to the non-deprived hemisphere with MD (Fig. S21).

**Figure 3.**
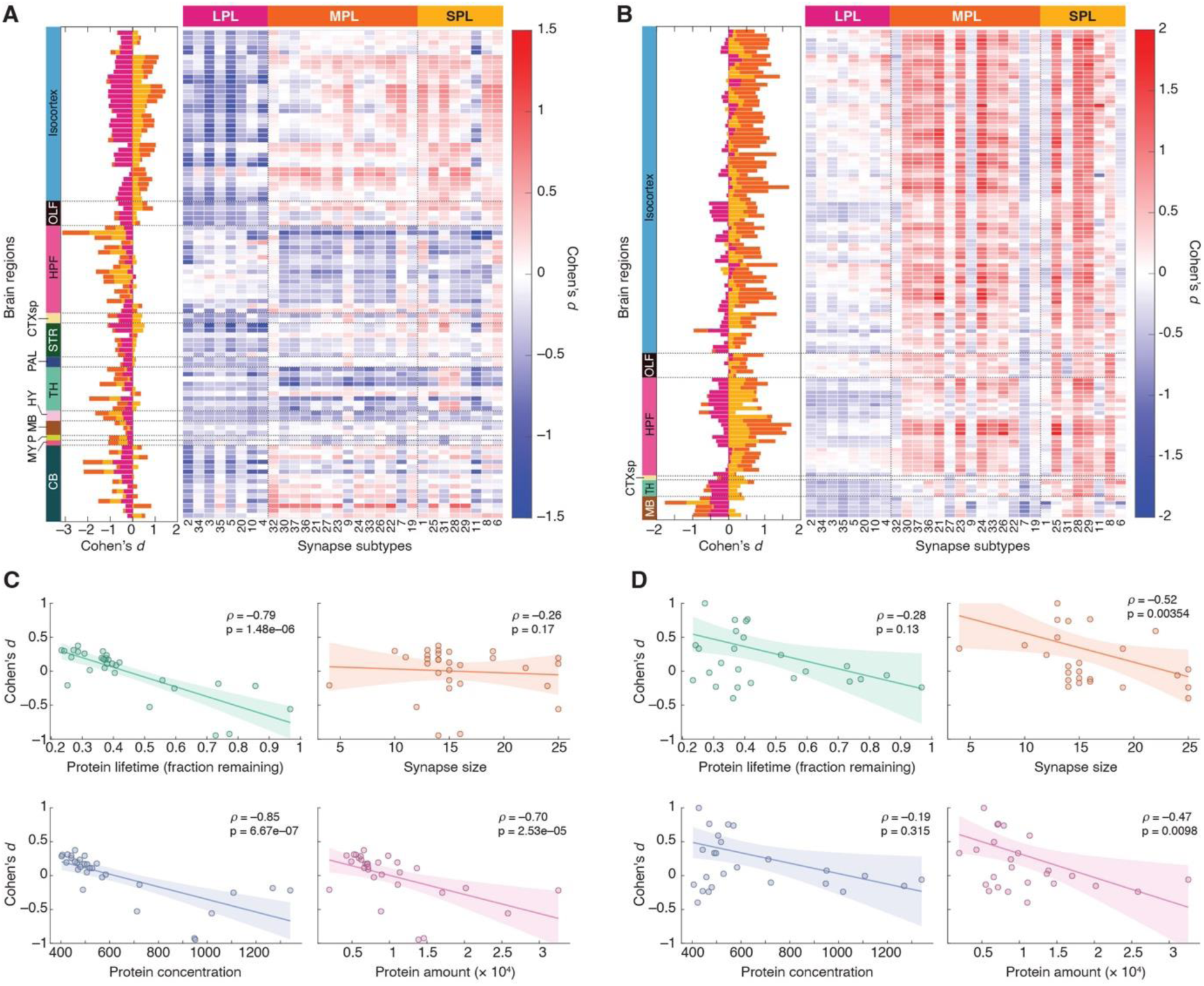
Experience-dependent plasticity selectively targets synapse subtypes according to their protein lifetime. (A, **B)** Regional effect sizes (Cohen’s *d*) for 30 PSD95⁺ synapse subtypes, ordered by protein lifetime, comparing developmental EE mice versus standard cage (SC) controls (A) and comparing monocular deprivation (MD) versus non-deprived controls (B). Stacked bar plots (left) show aggregate effect sizes for short (SPL, yellow; 8 shortest lifetime subtypes), medium (MPL, orange; 14 intermediate lifetime subtypes), and long (LPL, pink; 8 longest lifetime subtypes) lifetime groups. **(C, D)** Scatterplots showing the relationship between subtype effect size and protein turnover or structural features across 30 PSD95⁺ subtypes in the isocortex. Protein lifetime shows fraction remaining after 7 days; synapse size shows area in pixels; protein concentration shows mean pixel intensity of PSD95⁺ puncta; total protein amount is summed pixel intensity of PSD95⁺ puncta. Spearman’s ρ and associated *p*-values are shown.

In addition to protein lifetime, synapse subtypes also vary in protein amount, protein concentration and synapse size ^16,17^. We asked which of these parameters correlates with the effects induced by EE and MD. EE-induced changes were strongly and negatively correlated with protein lifetime (p= 1.48 x 10^-6^), protein concentration (p= 6.67 x 10^-7^), and protein amount (p= 2.53 x 10^-5^) but not with synapse size (p= 0.17) (Fig. 3C). By contrast, MD-related effect sizes showed only weak associations, with moderate negative correlations to synapse size (p=3.54 x 10^-3^) and protein amount (p=9.8 x 10^-3^), but no significant relationship to protein lifetime or concentration (Fig. 3D). These findings suggest that EE targets synapses with stable, high PSD95 content proteomes for downscaling, whereas MD shifts medium-sized synapses with more dynamic protein content. Together, these results indicate that synapse proteostasis guides the reprogramming of synaptome architecture in response to experience.

### Remodelling of synaptome architecture aligns with the connectome

In addition to a role of synapses with different protein turnover rates, we hypothesized that the connectivity between brain regions might contribute to the spatial reorganisation of synaptome architecture in response to experience. We integrated our synaptome data with connectome data from the Allen Mouse Brain Connectivity Atlas ^40,41^. From tracer injections across 129 source regions, we extracted projection densities to 73 parasagittal or 91 coronal target sites, aligning these to our synaptome architecture regions (Fig. S23). We then computed Spearman’s correlations between projection density and either control subtype density (ρᴰ) or absolute effect size following enrichment or deprivation (ρᶜᵈ), applying FDR correction across all comparisons.

Hierarchical clustering of ρᴰ and ρᶜᵈ revealed coherent subtype groupings with shared projection profiles in both control mice and those subject to EE or MD. In control mice at all ages tested (1 to 12 months), LPL synapse subtypes had their strongest associations with sensory pathways, including visual, auditory, somatosensory and motor thalamo-cortical circuit regions (Fig. S24-S26). The change in abundance of Type 3 MPL and SPL subtypes with MD correlated with projection densities from the visual pathway, including the lateral geniculate nucleus and visual cortices (boxed area in Fig. 4A, Fig. S24). These subtype groupings also overlapped with those identified by PCA, t-distributed Stochastic Neighbour Embedding (t-SNE) and Uniform Manifold Approximation and Projection (UMAP), supporting the interpretation that experience targets pre-wired subsets of excitatory synapses embedded within the brain’s sensory circuitry (Fig. 2D, 4B,C and S27).

**Figure 4.**
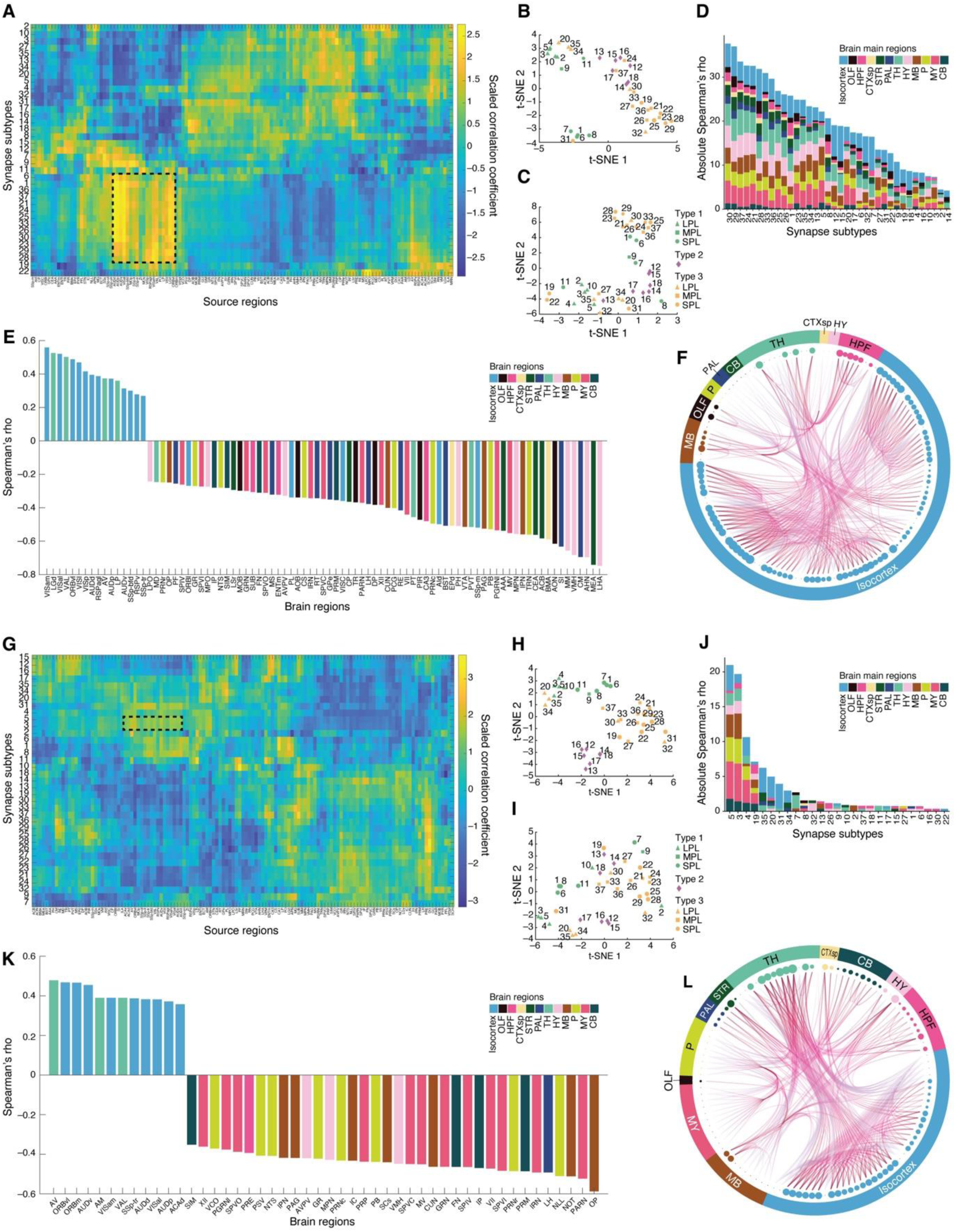
Synapse subtype remodelling under MD and EE aligns with mesoscale sensory connectivity. (A, **G)** Clustered heatmaps showing scaled Spearman correlation coefficients (ρCd) between synapse subtype effect size profiles (Cohen’s *d*) and projection densities from 129 source regions to 91 target regions, following monocular deprivation (MD; A) or to 73 target regions following juvenile environmental enrichment (EE; G). Rows represent 30 PSD95⁺ synapse subtypes; columns are anatomically defined source regions. Hierarchical clustering was applied to both axes using Pearson correlation distance. Boxed areas highlight positive correlations between clustered MD-affected (A) and EE-affected (G) subtypes in several thalamo-cortical sensory areas. **(B, H)** t-SNE projections of subtype density correlation profiles (ρD) in 1-month (B) and 3-month (H) control mice, revealing gradients that align with subtype protein composition and lifetime. Markers denote synapse type (see colour key) and protein lifetime class (see shape key). **(C, I)** t-SNE projections of subtype effect size correlation profiles (ρCd) under MD (C) and EE (I), revealing structured subtype groupings based on shared correlation signatures across source regions. Marker colours and shapes as B and H. **(D, J)** Stacked bar plots of significant absolute ρCd values (*p* < 0.019 for MD; *p* < 0.002 for EE; FDR-corrected) for each synapse subtype. Colours reflect source region groupings by anatomical domain. **(E, K)** Bar plots of significant source region correlations (ρCd) for representative subtypes: MD-responsive (subtype 29) and EE-responsive (subtype 5), highlighting key projection pathways associated with synapse remodelling. **(F, L)** Anatomically structured edge bundling graphs illustrating mesoscale projection networks linked to representative MD-responsive (subtype 29) and EE-responsive (subtype 5) subtypes. Nodes represent brain subregions coloured by anatomical domain; edges reflect 500 strongest projections from significantly correlated source regions. Node size denotes cumulative correlation magnitude (ρCd). Spearman’s ρ correlation profiles and t-SNE and UMAP embeddings are reported in the Supplementary Data file. For brain region abbreviations see supplementary material ‘Brain region list’.

Focusing on subtype 29, an SPL subtype upregulated by MD, we identified significant positive correlations with brainwide projections (p < 0.019, FDR-corrected; Fig.4D), with the strongest relationships from the visual pathway, including the lateral geniculate nucleus and anteromedial visual areas (Fig. 4E). A hierarchical bundling graph, limited to the 500 strongest projection pairs from significantly correlated source regions, highlighted the dense intra-cortical network underscoring these correlations (Fig. 4F). These findings show that the changes in synaptome architecture in response to MD are correlated with the inputs from the thalamo-cortical areas of the visual system.

EE revealed a distinct correlation profile from MD (Fig. 4A,G, S24 and S25). Among the few subtypes significantly correlated with projection densities, subtype 5 aligned with sensory thalamo-cortical projections, including those involved in spatial navigation, attention, and motor planning (Fig. 4G,J,K and S25). In both ρᴰ and ρᶜᵈ t-SNE and UMAP plots, LPL synapses were spatially grouped toward one end of the axis, indicating shared connectivity-correlation profiles for subtype control densities and EE effects (Fig. 4H,I and S27). A hierarchical bundling graph highlights the brainwide connections underscoring these correlation profiles (Fig. 4L). Together, these findings reveal that there is an intrinsic spatial relationship between the connectome and the synaptome architecture, which positions sensory pathways to drive changes in synapse populations in response to experience.

## Discussion

Our study demonstrates that environmental experience and sensory input drive widespread, selective remodelling of excitatory synapses across the brain in developing and mature mice. Using single-synapse resolution synaptome mapping of 37 varieties of excitatory synapses across more than 100 brain regions, we show that experience-induced changes are globally distributed yet type-and subtype-specific, forming regionally structured patterns in synaptome architecture. These patterns are distinct for different types of sensory experience and are correlated with large-scale connectivity and constrained by intrinsic synaptic proteostatic mechanisms. Importantly, we find that the same core mechanisms of synaptome reprogramming operate both during early developmental periods, including classical critical periods, and in the mature adult brain, indicating a conserved but flexible system for experience-dependent adaptation in populations of synapses across the lifespan.

The notion that experience is recorded in a distributed manner across many brain regions was first expounded by Semon^42^ and Lashley^43^ a century ago. In addition to Lashley’s pioneering experiments using cortical lesions, recent studies using reporters of activity-dependent genes^44–51^, calcium imaging^52,53^, multiunit recording with neuropixel probes^54–56^ and synapse protein imaging^57^, show that sensory stimuli and experience evoke highly distributed neural and even astrocyte^58^ changes. Taking these and our observations together, we propose that sensory experiences activate distributed neural networks, and that specific types and subtypes of synapses within neurons selectively respond to particular patterns of activity by remodeling their synapse proteomes. This results in synapse switching from one type or subtype to another, modifying the diversity of the synaptome. The net effect of this process is to leave a new synaptome architecture that is the record of the experience. Because the synapse types in our study are defined by the presence of PSD95 and/or SAP102, which are important regulators of synaptic strength and plasticity^59–62^, then the reprogrammed synaptome architecture would be expected to modify the physiological properties of networks and drive behavioural changes.

The distinct synaptome architectures induced by EE and MD indicate that different experiences leave contrasting molecular and spatial signatures within the synaptome architecture. These architectures emerge from the interplay of molecular and anatomical factors, including the molecular composition and proteostatic properties of synapses and their inter-regional connectivity. Storing representations in distributed synaptome changes would endow the brain with enormous capacity, predicted to scale exponentially with brain size.

Selective modification of subsets of synapses within a diverse population offers a new perspective on the classical model of synaptic plasticity, which emphasises changes in synaptic strength with experience. Our findings suggest a population-selection model of adaptation, analogous to processes in evolutionary biology, genetics and immunology. In evolution, selective pressures act on genetically diverse populations to shift allele frequencies^63^; in immunity, antigens select from a vast repertoire of pre-programmed lymphocytes, driving clonal expansion^64^. Similarly, we find that in the brain, environmental exposure acts on molecularly diverse synapses, altering the frequencies of synapse types and subtypes. Unlike genetic selection, which unfolds across generations, synaptic and immunological selection mechanisms enable rapid adaptation within the lifespan of an individual.

This synapse-selection and adaptation model has implications for understanding how experience and genetic factors interact. Synaptome architecture is established during development by a hierarchical cascade of genome-encoded molecular mechanisms, including transcriptional, translational, and post-translational processes that control the synthesis and spatial distribution of protein complexes across diverse synapse types in the dendritic arbor^16^. Supporting the central role of genetic mechanisms in the initial programming of the synaptome architecture, we have previously shown that autism-and schizophrenia-relevant mutations in transcription factors^22^ and synaptic proteins^16,17^ cause large-scale modification of synaptome architecture. Moreover, synaptome changes during development, ageing^17,21^ and after sleep deprivation^23^ indicate that diverse intrinsic and extrinsic influences converge on the synaptome as a common substrate. Thus, the spatial organisation of molecularly heterogeneous excitatory synapses into a dynamic synaptome architecture constitutes a flexible and versatile system through which genetic and environmental influences converge.

### A nexus between the synaptome, connectome and proteostasis

The distributed and selective nature of synaptome remodelling led us to explore the potential roles of synapse proteome stability (proteostasis) and the organization of the connectome. We found that the synapse subtypes most affected by EE were characterized by long PSD95 protein lifetimes and high protein concentrations, suggesting that enrichment selectively downscales stable, proteome-rich synapses. This aligns with a recent pixel-based analysis of PSD95 protein, showing faster turnover rates in a small group of adult mice housed in EE for two weeks compared to SC controls^57^. By contrast, MD preferentially increased subtypes with medium-or short-lived proteins. This suggests that the synaptic adaptation to experience and sensory input is dependent on the intrinsic (cell-autonomous) proteome stability, adaptability and molecular composition.

Although proteostatic mechanisms alone could achieve selective synapse modification, we found that the spatial distribution of LPL and SPL synapses was anatomically structured within the connectome, with LPL subtypes showing their strongest associations with sensory pathways. When we examined the subtypes upregulated in response to MD, we found the regions receiving strong thalamo-cortical visual inputs, such as the lateral geniculate nucleus and visual cortices, drove changes in SPL and MPL subtypes. These MD-induced synaptome changes were more extensively correlated with projection strength than those triggered by EE, which showed an enhancement in existing, control projection-subtype relationships. This dichotomy might reflect the brain’s evolved sensitivity to the multimodal input in EE compared with the more extreme and unnatural manipulation of MD that drives changes in more specific sensory pathways. Furthermore, this contrast underscores the difference between modulating normal sensory experience and responding to its abrupt loss: whereas EE fine-tunes existing synaptic patterns, MD necessitates more radical reconfiguration, echoing known circuit-level rewiring in visual cortex following deprivation^65–69^. The capacity to partially decouple from typical projection-subtype alignments allows the engagement of alternate or latent circuits.

### Environmental enrichment and deprivation

A growing body of evidence demonstrates that early sensory, cognitive, and physical experiences shape brain development, influence cognitive resilience, and modulate vulnerability to neuropsychiatric disease across the lifespan^70–78^. Against this backdrop, EE paradigms in animal models have frequently been interpreted to support enhanced synaptogenesis as a cellular mechanism underlying experience-driven cognitive and behavioural benefits^28–34^. Contrary to the classical view, we observed no significant increase in total synapse density across the brain in young or mature animals. Instead, EE restructured the synaptome by downregulating specific subsets of Type 1 and Type 3 synapses across cortical, thalamic, and cerebellar regions. This fine-scale remodelling underscores that EE restructures existing populations of synapses rather than expanding excitatory connectivity, and that cognitive improvements associated with EE likely arise from selective reorganization of synapse populations rather than wholesale proliferation. Synaptome changes were enriched in the deep layers of the cortex and in sensorimotor and subcortical regions, consistent with known enrichment-induced behavioural gains in learning and sensorimotor coordination^24,79,80^. This reorganization was not limited to early development: adult mice exposed to one month of EE displayed overlapping subtype shifts, indicating that the capacity for synaptome reprogramming persists into maturity^81–83^, although the regional focus of plasticity shifted away from deep cortical layers.

The MD paradigm confirmed that modality-specific sensory deprivation can elicit global synaptome remodelling. MD led to brainwide increases in Type 3 synapses, notably in both contralateral and ipsilateral isocortex and hippocampus, as well as in auditory and somatosensory regions. Our findings counter earlier assumptions that MD-induced plasticity is confined to the visual system and instead reveal a cortex-wide redistribution of excitatory synapse subtypes. Consistent with our conclusions, recent evidence showing altered neuronal activity beyond the visual cortex in mice subjected to MD^66^ supports the idea that synaptome changes may underlie broader functional reorganization of brain connectivity.

In conclusion, our findings show that the molecular diversity of excitatory synapses provides a fundamental substrate for recording experience across the brain. Beyond identifying how experience leaves its imprint, the organisation of heterogeneous synapse populations into a dynamic synaptome architecture—shaped by experience, ageing and genetic variation—offers new insight into the mechanisms underlying psychiatric and neurological disorders. The spatial patterning of these synapses forms a flexible framework through which genetic and environmental influences converge, highlighting the potential of synaptome mapping to reveal how disease-associated alterations may be exacerbated or mitigated by environmental or pharmacological interventions.

## Acknowledgments

We thank D. Maizels for artwork, C. Davey for editing, and M. Hawrylycz for providing R code used to convert JSON-formatted connectome datasets.

## Author contributions

Conceptualization: SGNG, FS, HW

Methodology: HW, ÜG, SCG, BN, GV, ER, NHK, FS

Investigation: HW, ÜG, SCG, BN, FS Software: ZQ, DD, KY

Formal Analysis: HW Visualisation: HW

Writing - Original Draft: HW

Writing - Review & Editing: SGNG, HW Supervision: SGNG

Funding acquisition: SGNG

## Data and materials availability

All raw data are available from BioStudies (https://ftp.ebi.ac.uk/pub/databases/biostudies/S-BSST/147/S-BSST2147/Files/The-Synaptome-Experience-Project/). Software code is available on Git (https://git.ecdf.ed.ac.uk/ddominic/the-synaptome-experience-project/-/blob/main/README.md).

## Supplementary Material

## Methods

### Resource availability

#### Lead contact

Further information and requests for resources and reagents should be directed to and will be fulfilled by the lead contact, Seth G. N. Grant.

#### Materials availability

This study did not generate new unique reagents.

#### Data and code availability

All data have been deposited on BioStudies and are publicly available as of the date of publication. DOIs are listed in the key resources table. Code developed for synaptome mapping, and analyses outlined below are available in Git repositories cited in the key resources table. Any additional information required to re-analyse the data reported in this paper is available from the lead contact upon request.

### Animals

#### Mouse lines

Male and female mice used for the comparison of environmental enrichment (EE) and standard cage (SC) during development were of mixed background from crossing C57Bl/6 mouse lines expressing Psd95^eGFP/eGFP^;Sap102^mKO2/mKO2^, as described by Zhu and colleagues (2018), with a 129S5 strain expressing Glun1^TAP/TAP^ (Zhu et al., 2018; Frank et al., 2016). Male and female mice used for monocular deprivation (MD) and control groups were C57Bl/6 colonies expressing Psd95^eGFP/eGFP^;Sap102^mKO2/mKO2^. Female mice from both strains were used for the adult EE experiment. Animal experiments were carried out according to UK Home Office regulations and were authorised by the Edinburgh University Director of Biological Services and the Joint Biological Services of Cardiff University.

#### Monocular Deprivation

Mice were housed in standard caging in groups of 2-5 mice per cage. Mice were weaned at P21 and divided into male or female cages. Mice were assigned to undergo MD surgery to achieve balanced male-female ratios for MD and control groups. MD surgery entailed suturing of the right eye from P25-26 to P31. Anaesthesia was induced with 3% isoflurane in 100% oxygen and maintained at 2% isoflurane. An injection of Metacam (1 mg/kg subcutaneous) was given to alleviate post-operative pain. The lid margins of the right eye were trimmed, and the lids sutured shut with three mattress sutures. A topical antibiotic ointment, chloramphenicol, was applied before animals were allowed to recover and were returned to the home cage. MD and control mice were housed in the same standard cage environment before and after surgery. Animals were checked daily to ensure that the eye remained closed. MD and control mice were sacrificed at P31 by perfusion and fixed brains were stored in 30% sucrose solution (w/v in 1 × PBS) for up to 6 days at “ice box” temperature. Thirteen mice (7 female) were included in the MD group, and fourteen mice (7 female) in the non-deprived control group.

#### Environmental Enrichment

For the developmental enrichment study, mice were bred in-house to generate litters for enriched environment (EE) or standard condition (SC) housing. Pregnant dams were distributed 7 days before parturition into EE (n = 4 per cage) or SC (n = 2 per cage) cages to facilitate social enrichment. Litters remained undisturbed until postnatal day 5. At weaning (P19–P28), pups were sex-separated and housed according to their assigned condition. EE mice were housed in groups of 12–13 per sex-matched cage; SC mice were housed in groups of 2–7, across multiple sex-matched cages. Ear-notching enabled litter identification. Dams were removed at weaning. Mice were perfused between P81 and P96. Final sample sizes were EE n = 23 (12 female), SC n = 29 (15 female). For the adult enrichment study, a separate cohort of female C57BL/6J mice (aged 6–12 months) were randomly assigned to either enriched or standard housing for 1 month prior to perfusion. EE mice (n = 20) and SC mice (n = 20) were housed in same-sex groups under the conditions detailed below. Male mice were excluded due to the aggression typically observed upon late transfer into group-housed enriched environments.

EE mice were housed in Marlau cages (Viewpoint, Lyon, France) following established protocols *(45, 51)*. These cages provide a two-level structure including a living area with three running wheels, a red plastic shelter, and access to food through a maze that was reconfigured every two days from twelve preset designs. SC mice were housed in standard open-top cages (floor area: 501 cm²) with a cardboard shelter and clear plastic tunnel for handling. Both EE and SC cages received the same bedding and nesting material. Food and water were provided ad libitum, and all mice were kept under a 12 h light/dark cycle. Cages were cleaned weekly. Bedding and nests were partially transferred between old and new cages to maintain familiar olfactory cues. EE mice were temporarily transferred to standard cages during cage cleaning. All mice were handled using plastic tunnels, and handling was restricted to two experimenters to minimise variability.

### Sample preparation

#### Tissue Collection and Processing

Mice were deeply anaesthetised via intraperitoneal injection of sodium pentobarbital (20% w/v; Pentobarbitone Sodium, Animalcare; dose: 0.05–1.5 mL according to body size). Upon loss of pedal reflexes, mice underwent transcardial perfusion with 10 mL of phosphate-buffered saline (PBS, 1X; Oxoid), followed by 10 mL of 4% paraformaldehyde (PFA; Alfa Aesar) in PBS.

Brains were dissected and post-fixed in 4% PFA at 4 °C for 3–4 hours, then cryoprotected in 30% sucrose (w/v in PBS) at 4 °C for 3–6 days. Once equilibrated, brains were embedded in optimal cutting temperature (OCT) compound (CellPath) within cryomolds, snap-frozen in liquid nitrogen-cooled isopentane, and stored at −80 °C until sectioning.

#### Sectioning

Frozen brains were sectioned at 18 μm thickness using a cryostat. Sections were collected in either the coronal or parasagittal plane depending on the study. For the monocular deprivation experiment, coronal sections were collected from Bregma−3.07 mm to −3.27 mm (corresponding to images 85–86 of the Allen Mouse Brain Atlas). For synaptome mapping in enriched and standard cage mice, parasagittal sections were collected centred on the region corresponding to image 11 of the Allen Mouse Brain Atlas sagittal reference *(33)*. Two adjacent sections were mounted per slide (Superfrost Plus, Thermo Scientific), dried overnight at room temperature and then stored at −20 °C until imaging.

#### Mounting for Imaging

Slides were thawed at room temperature for 5 minutes before mounting. Sections were rehydrated in 1X PBS (Sigma-Aldrich) for 5 minutes in a light-protected humidity chamber. A volume of 12 μL of homemade Mowiol mounting medium (Calbiochem) was applied per section. Coverslips (13 mm or 18 mm × 1.5 mm; VWR International) were selected based on section orientation (coronal or parasagittal, respectively). Mounted slides were dried overnight in the dark at room temperature. For short-term use (<5 days), slides were stored at 4 °C; for longer-term storage, they were kept at −20 °C.

### Imaging and Data Acquisition

Whole-brain section imaging was performed using a Nikon Eclipse Ti2 inverted microscope equipped with a Yokogawa CSU-W1 spinning-disk confocal scanner and a Photometrics Prime BSI camera. Image acquisition was controlled via Nikon NIS-Elements software (v5.20.1–5.21.1). Each slide was first scanned at 4X magnification (NA 0.2) to acquire an overview of the brain sections and define coordinates for high-resolution imaging. Whole-section imaging was then performed at 100X magnification (NA 1.45, oil immersion), using automated tile scanning protocols with consistent acquisition settings across all experiments.

For high-magnification imaging of PSD95-eGFP and SAP102-mKO2, the 561 nm laser (SAP102-mKO2) was triggered first, followed by the 488 nm laser (PSD95-eGFP). Excitation was delivered through the L100 light path using emission filter 1. Laser power and exposure times were set to 2.33 mW for 200 ms (561 nm) and 1.06 mW for 120 ms (488 nm), respectively. Aperture was set to 1, and imaging was performed in 12-bit with Correlated Multi-Sampling (CMS) gain.

For low magnification 4X overview imaging, laser powers were reduced to minimise photobleaching: 0.61 mW for 561 nm (200 ms) and 0.71 mW for 488 nm (120 ms), with aperture set to 7. The same light path (L100), emission filter (1), gain (CMS), and 12-bit resolution were used as in the high-magnification protocol.

Each whole-brain scan contained over 10,000 tiles. Autofocus correction was performed every 400 tiles using a z-stack of 41 optical sections (0.5 µm step size). Final z-positions were calculated from the midpoint intensity value of the stack to account for focal drift along the z-plane. Autofocus points were manually selected in regions of high synaptic density.

For the adult EE dataset, we used a subsampled variant of the Synaptome mapping imaging protocol to increase throughput. Instead of contiguous rectangular tiling, high-magnification (100×) imaging was performed on a systematic sparse-sampling grid: fields of view were acquired at regular offsets in X and Y with a centre-to-centre step equal to 1.5× the field of view, yielding non-overlapping images separated by half a field. Polygonal ROIs were defined from the 4× overview as above, and the grid was applied uniformly within each ROI. All other optical and acquisition parameters and the autofocus procedure matched the whole-section protocol, providing a balanced compromise between mapping speed and spatial precision.

All scans were acquired using automated JOBS routines in NIS-Elements, incorporating the optical configuration settings, autofocus correction, polygonal ROI assignment, and coordinate interpolation. Imaging settings were validated using saturation and signal-to-noise ratio (SNR) analyses. SNR was optimised using TrackMate in ImageJ (v2.0.0) with custom MATLAB scripts (available on request).

### Image Processing and Brain Region Delineation

Following whole-brain section imaging, raw image files were processed using Nikon NIS-Elements Advanced Research software (v5.21). Image metadata and individual 856×812-pixel tiles were extracted for each section. To evaluate image quality and anatomical positioning, upscaled overview montages were generated using custom MATLAB scripts (R2020b–R2022a; scripts available upon request). These montages were reconstructed using the extracted tiles and metadata, enabling visual inspection for imaging artefacts, tissue damage, and scan alignment along the anteroposterior and mediolateral axes. Scans with extensive out-of-focus areas—primarily due to drift in the nosepiece z-plane—were excluded and repeated using adjacent sections.

Montage images from accepted scans were used for manual delineation of brain regions, guided by the Allen Mouse Brain Atlas *(33)*. One imaged section was used per mouse for delineation and subsequent analysis. Delineations were performed in ImageJ (v2.14.0) using the polygon ROI tool and ROI Manager. Only visually distinguishable subregions corresponding to the reference atlas image were included. For manual delineations, synaptic marker expression (PSD95-eGFP and SAP102-mKO2), cell body density, and laminar organisation were used in conjunction with atlas references to guide boundary placement. In the monocular deprivation study, additional reference was made to cortical layer 4 morphology to distinguish visual subfields (Antonini et al., 1999). All delineated regions used in analysis are listed in the supplementary materials (Fig. S27 and S28). While anatomical landmarks and synaptic marker patterns were used to approximate regional boundaries, it is acknowledged that these may not precisely reflect functional or synapse-specific zones.

Manual brain region delineation was performed for the MD and developmental EE studies. For the adult EE study, an automated delineation tool developed in-house was employed, which had been trained using the manual delineations from existing datasets *(63)*. Automated delineations of each section underwent visual QC. When a boundary showed a clear deviation from the atlas or from our manual delineation criteria, the affected ROI mask PNGs were imported into napari (v0.5.5) as label layers and corrected with pixel-level paint/erase tools to realign boundaries to cytoarchitectonic landmarks and synaptic marker patterns. Edits were sparse and local: many sections required no changes, while others required adjustments to fewer than 15 subregions, most commonly laminar boundaries in PTLp and cerebellar lobules. Corrected masks were exported as updated PNGs and used for downstream ROI assignment and quantification.

### Synaptome Mapping: Quantification of synapse parameters

#### Punctum Detection

To identify PSD95-eGFP and SAP102-mKO2 synaptic puncta, we used a custom machine learning-based detection algorithm adapted from previously validated synaptome mapping pipelines (Zhu et al., 2018; Cizeron et al., 2020). Detection was performed independently for the PSD95 and SAP102 channels. Fluorescently tagged puncta were detected using a multi-scale, multi-orientation feature extraction framework combined with an ensemble-trained neural network classifier. To adapt the synaptome mapping detection algorithm for use with the Nikon Ti2 Eclipse spinning-disk confocal system, we generated a new training dataset comprising z-stacks (5 planes, 150 nm spacing) from parasagittal brain sections of *Psd95*^eGFP/eGFP^; *Sap10*2^mKO2/mKO2^ mice aged P7, P14, 3 months, and 1 year (n = 3 per age group). Randomised 128 × 128 pixel sub-tiles (n = 1000) were extracted and manually annotated using the ImageJ cell counter plugin by five independent raters blinded to study aims. Puncta were defined as bright, round or elliptical spots, and annotations were minimally guided to reduce selection bias. These manual annotations were used as training labels for the classifier. Candidate puncta were extracted using local image features, then classified as true synaptic puncta or background based on their feature vectors. Final outputs included the coordinates and extracted properties for each detected punctum. Code and trained classifiers are available on GitHub (see Key Resources Table).

### Punctum Segmentation and Parameter Extraction

Following detection, each synaptic punctum was segmented using a local thresholding approach based on 20% of its peak intensity, analogous to full width at half maximum (FWHM). This allowed separation of size and intensity measurements. A marker-controlled watershed algorithm was applied to resolve overlapping puncta. Puncta smaller than 4 pixels were excluded. For each segmented punctum, six parameters were calculated: mean intensity, area, circularity, aspect ratio, skewness, and kurtosis. Shape descriptors quantified the asymmetry, peakedness, and roundness of each punctum. Parameter extraction was performed using MATLAB’s *regionprops* function and custom scripts. This step enabled downstream synapse classification based on both molecular composition and morphological features.

#### Punctum Colocalisation

To determine which puncta co-expressed PSD95 and SAP102 at the same synapse, we performed a spatial colocalisation analysis based on Euclidean distance between punctum centroids (Zhu et al., 2018). Pairs separated by ≤500 nm were classified as colocalised. This threshold was selected based on both synaptic ultrastructure (Sheng & Kim, 2011) and dual-peaked histograms of punctum distances. Due to the diffraction limit, only one punctum per protein is expected per synapse, so a one-to-one matching constraint was imposed and modelled as a linear assignment problem, solved using the Hungarian algorithm. Colocalisation was performed after independent detection and segmentation of both channels.

#### Punctum Classification

To capture synapse diversity, each punctum was classified into one of 37 previously defined synapse subtypes (Zhu et al., 2018) based on size, intensity, and shape parameters. Originally derived using unsupervised weighted ensemble clustering (WEC) on Andor microscope data, this classification system was adapted for use with Nikon-imaged data via transfer learning: the original classifier structure was preserved, but certain model layers were fine-tuned on Nikon-imaged puncta using densely connected convolutional networks (DenseNet). Three separate classifiers were trained for PSD-95+ only, SAP-102+ only, and colocalised PSD-95+/SAP-102+ puncta using puncta-specific features: Mean Intensity, area, skewness, kurtosis, circularity and aspect ratio. Classifiers were implemented as Compact Classification ECOC models in MATLAB. All classifiers are available on GitHub (see Key Resources Table). Subtype labels and confidence scores were assigned using the built-in MATLAB ‘predict’ function. Classifier outputs were used in downstream brain mapping analyses.

#### Brain Mapping

Following detection, colocalisation, and classification, synaptic parameters were aggregated to anatomical brain regions. First, punctum-level features were averaged within 4 × 4 subtiles per image tile. These subtile-level summaries were then mapped to anatomical subregions delineated using the Allen Mouse Brain Atlas *(33)*. Regional values (e.g., intensity, size, shape, subtype density) were computed by averaging across all subtiles within each delineated mask. Puncta exceeding 140 pixels in size or with classification confidence <10% were excluded. Puncta on tile edges were also excluded to reduce border artefacts. Final outputs for each region included average values of synaptic parameters and subtype compositions, exported as Excel spreadsheets for downstream analysis.

## Statistical Analyses

### Data Handling

Raw TIFF images, metadata, and delineated ROI files were stored and archived on institutional secure servers. Processed outputs from the SYNMAP and TrackMate pipelines—including punctum parameters, colocalisation metrics, and region-level summary values—were saved in Excel format and analysed using MATLAB (R2020b– R2022a) and R (v4.4.0). For each animal, punctum and synapse subtype data were aggregated by anatomical subregion *(27)*. Group *n* values reported in the main text reflect the number of animals unless otherwise indicated. In some cases, subregions do not include data from all animals, either because the small size and anatomical delineation of certain areas precluded averaging in the SYNMAP pipeline, or because those regions were excluded during delineation. Subregion-level *n* are provided in the Supplementary Data. Bayesian statistical analysis was performed using adapted open-source MATLAB and R code (see Key Resources Table). Figures were generated using MATLAB, Rstudio and a brain heatmap plotting tool developed in house (see Key Resources Table). Due to the granularity of the Allen Mouse Brain Atlas reference maps, the primary visual cortex area of the coronal brain heatmap tool shows only binocular zone data. Both binocular and monocular zone data are plotted in the simple heatmaps.

### Bayesian Analysis

To assess experience-and sex-dependent differences in synapse parameters across brain subregions, we implemented robust Bayesian estimation models adapted from Kruschke’s hierarchical framework as per previous synaptome mapping publications (Zhu et al., 2018; Cizeron et al., 2020; Bulovaite et al., 2022; Koukaroudi et al., 2024; Tomas-Roca et al., 2022; Kruschke, 2013, 2014). These models were applied separately for each synapse parameter, across multiple brain subregions, for three experimental conditions: MD, developmental EE, and adult EE. All comparisons were performed against matched control groups.

### Two-Group Comparisons

For comparisons involving only experimental versus control animals (e.g. adult EE), Bayesian estimation was used as a principled alternative to t-tests. Each dataset was modelled using a Student’s t-distribution with group-specific means (μ₁, μ₂) and standard deviations (σ₁, σ₂), and a shared degrees of freedom parameter (ν) to capture potential outliers. Priors for μ were normal distributions centered on the pooled data mean, while σ values were assigned gamma priors based on empirical variance. The ν parameter followed a truncated exponential prior (rate = 1/30, lower bound = 1). Posterior samples were drawn using MCMC via matjags (MATLAB) using 4 independent chains, 2,000 burn-in iterations per chain, and thinning by 5. We retained 3,750 post–burn-in samples per chain (15,000 posterior draws total) for each monitored parameter Posterior distributions of μ₁–μ₂ were used to evaluate group differences.

### Factorial Comparisons

To assess interaction effects between sex and condition (e.g. MD or developmental EE), we used a Bayesian two-way ANOVA structure implemented in JAGS via the runjags package in R. Observed values yₖ were modelled as yₖ ∼ t(μₖ, σ²₍ⱼ₁,ⱼ₂₎, ν), where μₖ comprised additive and interaction terms corresponding to sex, condition, and their interaction. Each effect had a zero-centered normal prior with empirical scaling, and group-specific variances were modelled hierarchically with gamma hyperpriors. All posterior draws were reparameterized post hoc into sum-to-zero contrasts (b₀, b₁, b₂, b₁b₂) for interpretation of main and interaction effects. Models were fit in JAGS via runjags (R) using MCMC with 4 chains, 2,000 burn-in iterations per chain, thinning by 5, and 3,750 retained post–burn-in samples per chain (15,000 posterior draws total).

### Posterior Inference and Significance Estimation

We used the posterior distribution of the group difference (μ₁–μ₂ or specified contrasts) to assess evidence for group differences as per previous synaptome mapping publications (Zhu et al., 2018; Cizeron et al., 2020; Bulovaite et al., 2022; Koukaroudi et al., 2024; Tomas-Roca et al., 2022). A Region of Practical Equivalence (ROPE) was defined as ±⅓σ around zero, where σ denotes the posterior standard deviation of μ₁–μ₂. To approximate a p-value, the width of the Highest Density Interval (HDI) was iteratively narrowed from the full posterior until the HDI excluded the ROPE. The posterior mass outside this HDI was reported as a p-like metric reflecting credible evidence against the null hypothesis. These p-values were calculated independently for each brain subregion and synapse subtype parameter.

### Multiple Comparisons Correction

For each synapse subtype, Benjamini-Hochberg (BH) correction was applied across all subregions (i.e., 97 (adult EE), 101 (developmental EE) and 256 (MD)) to control the false discovery rate (FDR = 0.05) as per previous synaptome mapping publications (Zhu et al., 2018; Cizeron et al., 2020; Bulovaite et al., 2022; Koukaroudi et al., 2024; Tomas-Roca et al., 2022). This method preserves power to detect localized, biologically meaningful differences while limiting false positives. We note that BH correction assumes independence, which may not hold across anatomically connected regions. Nevertheless, this procedure offers a conservative benchmark for significance. All custom model code, and posterior processing functions are available on GitHub (see Key Resources Table).

### Derived Metrics and Statistical Calculations Synapse Type and Total Synapse Densities

For each mouse and brain subregion, synapse densities were computed by summing the subtype-level densities from the synaptome mapping pipeline. Synapse type densities were computed by summing the densities of their corresponding subtypes: Type 1 (PSD95-only, subtypes 1–11), Type 2 (SAP102-only, subtypes 12–18), Type 3 (colocalised, subtypes 19–37), and total synapse density (subtypes 1–37).

### Effect Size Estimation (Cohen’s d)

Cohen’s d was calculated to estimate the effect size between groups using the pooled standard deviation formula:

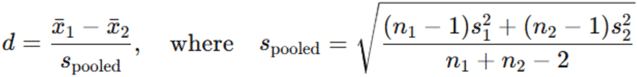

Here, x_1_ and x_2_ are group means, s_1_ and s_2_ are standard deviations, and n_1_, n_2_ are group sample sizes.

### Cortical Area Normalisation for Heatmaps

To visualize relative changes across cortical laminae, each subtype-layer effect size within a cortical area was normalized by dividing by the maximum observed effect size across all layers and subtypes of that area.

### Inter-Hemisphere Difference and Normalised Contralateral Indices

To quantify lateralized responses to monocular deprivation (MD), the following metrics were computed for each punctum and subtype parameter:

Asymmetry Index:

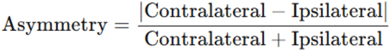

Normalised Contralateral Index:

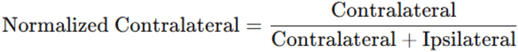

Both metrics were computed per individual and averaged at the group level. The asymmetry index captures magnitude of lateralisation, while the normalised index reflects directional bias.

### Similarity matrices

To compare which subregions and subtypes were similarly affected by environmental enrichment, monocular deprivation, or between enrichment paradigms, Pearson’s correlation similarity matrices were generated for Cohen’s d effect size subregion x subtype matrices.

### PCA-clustering analysis

Cohen’s d effect size values were computed for each of 37 synapse subtypes across all anatomically defined brain subregions in each experimental condition (MD vs. non-deprived control; EE vs. SC control). Prior to analysis, variables were centred and scaled. Assumptions regarding normality, multicollinearity, and linearity were assessed; robust methods were used where appropriate. Robust PCA was performed using the PcaHubert() function (R package rrcov, v1.6), which reduces sensitivity to outliers and non-Gaussian distributions. The number of retained principal components was determined by visual inspection of scree plots and bootstrapped confidence intervals (1,000 resamples). PCA scores were subjected to agglomerative hierarchical clustering using Ward’s method. The optimal number of clusters was identified via the NbClust consensus method. Cluster robustness was assessed using Jaccard stability indices (bootstrap resampling, 1,000 iterations), and Kruskal–Wallis tests were used to assess significant separation between clusters along principal components. All analyses were conducted in R (v4.3.1).

### Spearman’s correlation of subtype parameters

To examine how synapse subtype traits relate to their responsiveness to environmental enrichment, Spearman’s rank correlations were computed between Cohen’s d effect sizes (calculated for each PSD95⁺ subtype in the isocortex) and subtype-specific parameters: protein lifetime, protein concentration, protein amount, and synapse size. Correlations were computed using Spearman’s rank method (ρ) via the corr function in MATLAB (R2022a). Subtype lifetimes were sourced from Bulovaite et al. (2022). Protein concentration was estimated using the mean pixel intensity of PSD95 puncta, and punctum size was defined by the pixel area. Protein amount was calculated as the product of mean intensity and area. All parameters were derived from the synaptome mapping pipeline.

### Connectivity correlation analysis

To investigate whether mesoscale axonal connectivity constrains synaptome organisation and plasticity, we correlated synapse subtype metrics with anatomical projection strengths obtained from the Allen Mouse Brain Connectivity Atlas *(32)*. Projection data were sourced from 129 tracer injection experiments in wild-type mice, each reporting projection densities from a single source region to multiple annotated brain targets. A metadata.csv file and associated per-experiment XML files were downloaded manually from the Allen portal using a custom Python script. XML data were converted to readable format using an R script provided by the Allen Institute.

Projection density values, injection volumes, experiment labels, and region labels from each injection experiment were extracted using custom MATLAB scripts. Projection densities were normalised by the injection volume per experiment. Where multiple experiments targeted the same source region, values were averaged to obtain a representative profile.

To align our synaptome-mapped brain subregions with available connectome targets, we manually matched delineated anatomical regions to Allen-defined targets using the Allen Reference Atlas *(33)*. Regional groupings (e.g., cortical layers, visual subfields) were averaged where needed to harmonise naming across datasets. This mapping step involved averaging synapse metrics across some adjacent subregions where the connectome dataset did not distinguish between them. Separate connectome matrices were produced for the monocular deprivation (MD) dataset (coronal sections; 91 target regions) and the environmental enrichment (EE) dataset (sagittal sections; 73 target regions) due to differences in imaging coverage.

For each experiment, baseline synapse subtype densities (mean of control animals) and absolute Cohen’s d effect sizes were compiled into 37 × N matrices, where N is the number of matched target regions per dataset. These matrices were then correlated against a normalised 129 × N matrix of projection densities using Spearman’s rank correlation (MATLAB corr function,’Type’,’Spearman’). P-values were derived using MATLAB’s default large-sample approximation for Spearman’s rho, and multiple comparisons were corrected using the Benjamini–Hochberg method (built-in MATLAB function). Correlations with adjusted p-values below the critical threshold were classed as significant.

### Clustering heatmap

To visualise shared projection associations among synapse subtypes, we clustered correlation matrices relating projection strength to either baseline subtype density (ρᴰ) or experience-induced effect size (ρᶜᵈ). Each matrix (subtypes × source regions) was z-scored across projection regions to normalise subtype profiles. Hierarchical clustering was applied to both rows and columns using Pearson correlation distance and average linkage. Optimal leaf ordering was used to enhance cluster visualisation. Heatmaps were generated in MATLAB using the reordered matrices and labelled axes to identify synapse subtypes with similar projection correlation patterns.

### T-SNE and UMAP analysis

To visualise projection associations across synapse subtypes, we applied dimensionality reduction to matrices of Spearman’s rank correlation coefficients relating projection strength to either baseline subtype density (ρᴰ) or enrichment-or deprivation-induced effect sizes (ρᶜᵈ). Each region was standardised across synapse populations using the scale() function in R (z-scoring to zero mean and unit variance), so that all regions contributed comparably to the analysis. We used both t-distributed Stochastic Neighbour Embedding (t-SNE) from the Rtsne package in R and Uniform Manifold Approximation and Projection (UMAP) from the uwot package in R, using euclidean-based distance metrics. For t-SNE, perplexity values from 5 to 12 were tested; for UMAP, the number of nearest neighbours varied from 5 to 12. Parameter optimisation was guided by Mantel correlations between high-and low-dimensional distance matrices, reflecting global structure preservation. To evaluate local fidelity, we calculated trustworthiness and k-nearest neighbour overlap (k = 5) between high-dimensional and embedded spaces. Embedding stability was assessed by performing ten replicate runs and measuring alignment consistency via Procrustes analysis. Final embeddings used in figures were generated using a perplexity of 12 for t-SNE, and 10 (1M/MD) or 8 (3M/EE) nearest neighbours for UMAP. For t-SNE, the most representative embedding was selected as the run with the lowest average Procrustes residual across replicates, and this solution was used in final MATLAB visualisations.

### Anatomically Structured Edge Bundling of Synapse–Projection Relationships

To visualise how experience-dependent synapse plasticity integrates with the brain’s mesoscale connectivity, we constructed a hierarchical bundling graph in a circular dendrogram layout. Nodes represented brain subregions, grouped into major anatomical domains. Edges represented projection strength between source and target regions based on connectivity data from the Allen Mouse Brain Connectivity Atlas. Projection strength and Spearman’s correlations were combined to generate a weighted edge list. Only statistically significant connections (p ≤ critical p) were retained, and the top 500 strongest source–target pairs were visualised. A custom MATLAB pipeline processed, filtered, and ranked edges based on projection strength and correlation magnitude (see Key Resources Table). The final node parameter values (summed correlation strengths) were computed per subregion for available target region data. Graph layout and plotting were performed in R using the ggraph and igraph packages. The hierarchy was defined by parent–child relationships between anatomical domains and subregions, with an artificial central “origin” node anchoring all domains. Subregion nodes were coloured by anatomical domain using a custom colour map.

## Key Resources Table

| Chemicals, peptides, and recombinant proteins |  |  |
| --- | --- | --- |
| OCT embedding matrix | Cell Path | #KMA-0100-00A |
| Deposited data |  |  |
| Allen mouse brain reference atlas | Allen Reference Atlas – Mouse Brain | http://mouse.brain-map.org/static/atlas |
| Mesoscale structural connectome | Allen Mouse Brain Connectivity Atlas | http://connectivity.brain-map.org/ |
| Cohen's d effect sizes, p-values, similarity matrix, synapse subtype intensity, size, protein amount and lifetime values, synaptome-connectome correlation matrices, and t-SNE/UMAP embeddings | This paper OR BioStudies | Supplementary Data OR<br>https://doi.org/10.6019/S-BSST2147<br>https://ftp.ebi.ac.uk/pub/databases/biostudies/S-BSST/147/S-BSST2147/Files/The-Synaptome-Experience-Project/ |
| Synaptome data with individual synapse parameters and subtypes across all mouse brain sections | BioStudies | https://doi.org/10.6019/S-BSST2147<br>https://ftp.ebi.ac.uk/pub/databases/biostudies/S-BSST/147/S-BSST2147/Files/The-Synaptome-Experience-Project/ |
| Brain region list with abbreviations | This paper | Supplementary Material |
| Posterior distributions (MCMC draws) | Git | https://git.ecdf.ed.ac.uk/ddominic/the-synaptome-experience-project/-/blob/main/README.md |
| PCA scores and loadings | Git | https://git.ecdf.ed.ac.uk/ddominic/the-synaptome-experience-project/-/blob/main/README.md |
| Projection matrix (source × target, normalised) | Git | https://git.ecdf.ed.ac.uk/ddominic/the-synaptome-experience-project/-/blob/main/README.md |
| Synaptome-connectome region mapping table | Git | https://git.ecdf.ed.ac.uk/ddominic/the-synaptome-experience-project/-/blob/main/README.md |
| Edge list for bundled graph (source, target, weight, correlation) | Git | https://git.ecdf.ed.ac.uk/ddominic/the-synaptome-experience-project/-/blob/main/README.md |
| Software and Code |  |  |
| MATLAB | MathWorks | https://uk.mathworks.com/?s_tid=gn_logo |
| R for statistical computing | R projects | https://www.r-project.org/ |
| ImageJ | NIH | https://imagej.net/ij/ |
| Bayesian estimation | Git and GitHub | https://git.ecdf.ed.ac.uk/ddominic/the-synaptome-experience-project<br>https://github.com/NilsWinter/matlab-bayesian-estimation<br>https://github.com/boboppie/kruschke-doing_bayesian_data_analysis/tree/master/2e |
| Just Another Gibbs Sampler | SourceForge | https://sourceforge.net/projects/mcmc-jags/files/ |
| Brain map plotting tool | Git | https://git.ecdf.ed.ac.uk/ddominic/the-synaptome-experience-project/-/blob/main/README.md |
| Synaptome Mapping pipeline | Git | https://git.ecdf.ed.ac.uk/ddominic/the-synaptome-experience-project/-/blob/main/README.md |
| PCA + hierarchical clustering + NbClust | Git | https://git.ecdf.ed.ac.uk/ddominic/the-synaptome-experience-project/-/blob/main/README.md |
| Connectivity correlation | Git | https://git.ecdf.ed.ac.uk/ddominic/the-synaptome-experience-project/-/blob/main/README.md |
| Dimensionality reduction (t-SNE, UMAP) | Git | https://git.ecdf.ed.ac.uk/ddominic/the-synaptome-experience-project/-/blob/main/README.md |
| Edge bundling visualisation (circular dendrogram) | Git | https://git.ecdf.ed.ac.uk/ddominic/the-synaptome-experience-project/-/blob/main/README.md |
| Allen API parsing and XML preprocessing | Git | https://git.ecdf.ed.ac.uk/ddominic/the-synaptome-experience-project/-/blob/main/README.md |
| z-scoring and hierarchical clustering (heatmaps) | Git | https://git.ecdf.ed.ac.uk/ddominic/the-synaptome-experience-project/-/blob/main/README.md |
| **Other** |  |  |
| Paper resources | Git | https://git-pages.ecdf.ed.ac.uk/the-synaptome-experience-project-289b92/ |

---

**Fig. S1).**
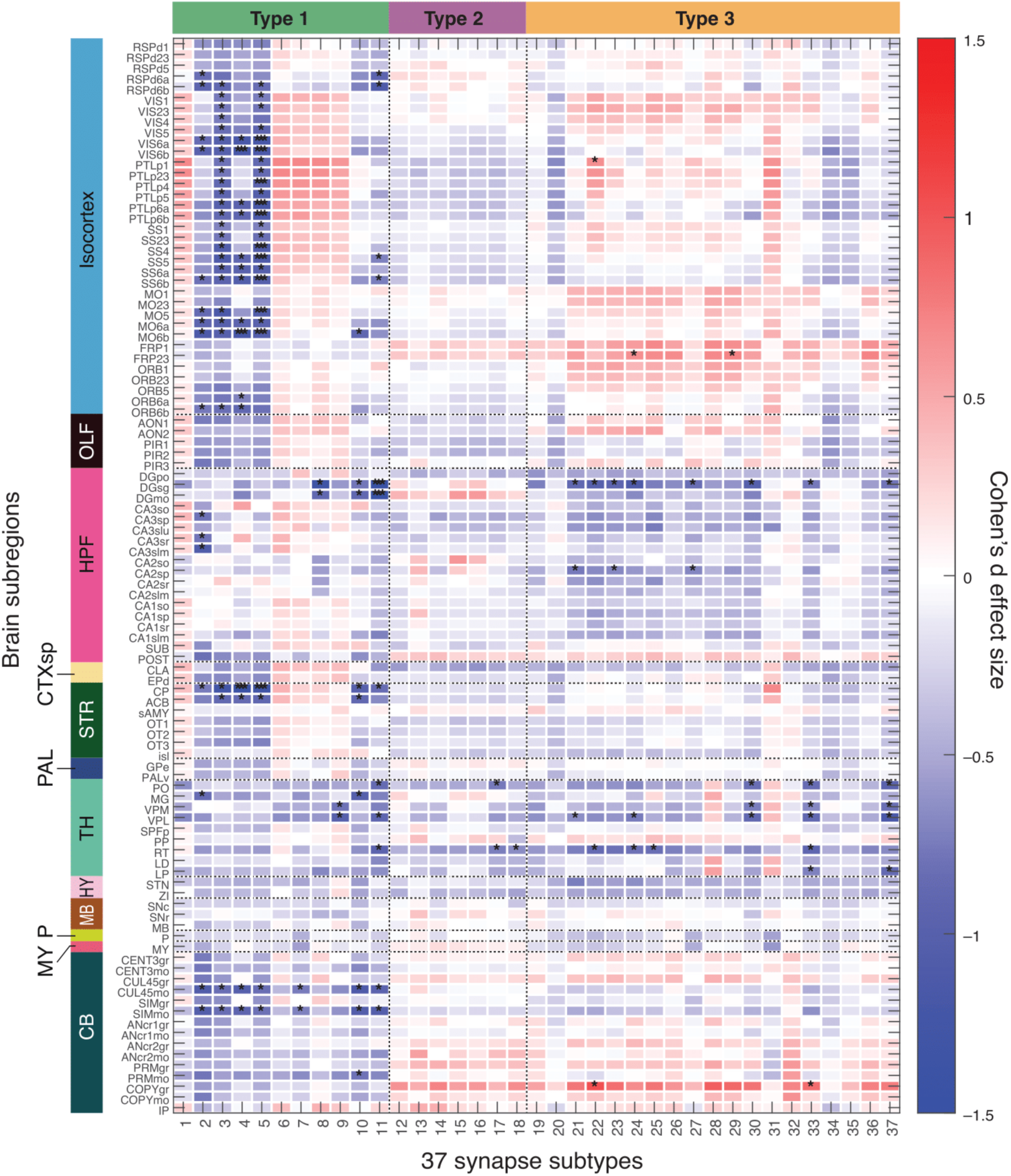
Synapse subtype density differences across all brain subregions between developmental EE and SC mice. Heatmap shows Cohen’s *d* effect size for 37 synapse subtypes; red indicates higher density in EE mice. Asterisks denote statistical significance (*p < 0.05; ***p < critical p after Benjamini–Hochberg correction applied to Bayesian posterior probabilities). Significance after correction was found for subtypes 4, 5, and 11 (p < 0.0014, 0.0061, 0.00047, respectively). For brain region abbreviations see supplementary material ‘Brain region list’.

**Fig. S2).**
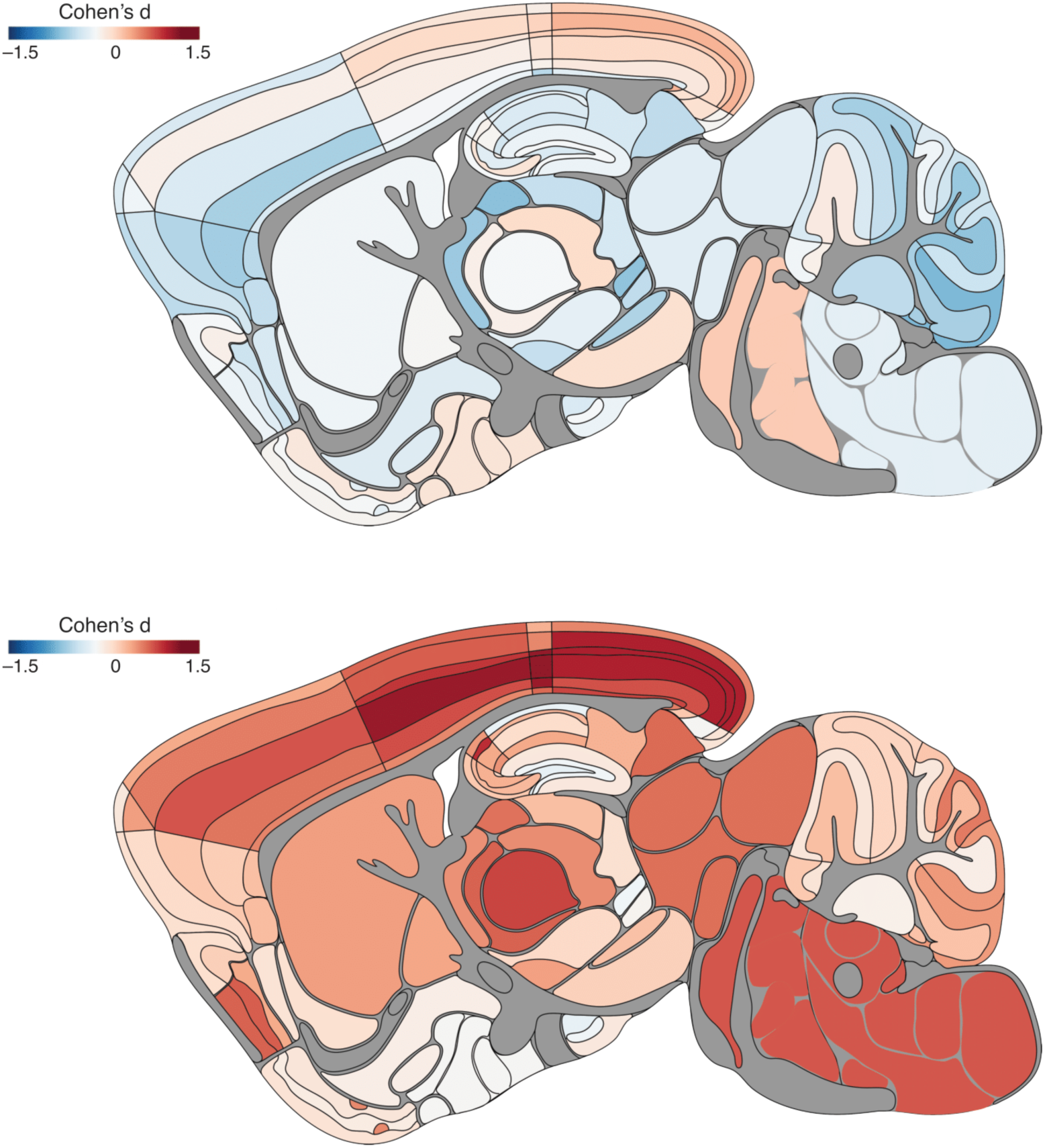
Parasagittal brain maps comparing total synapse density between female and male mice in 3-month SC (top) and developmental EE (bottom) groups with Cohen’s *d* effect size. Red indicates higher density in females. See Fig. S29 for region key.

**Fig. S3).**
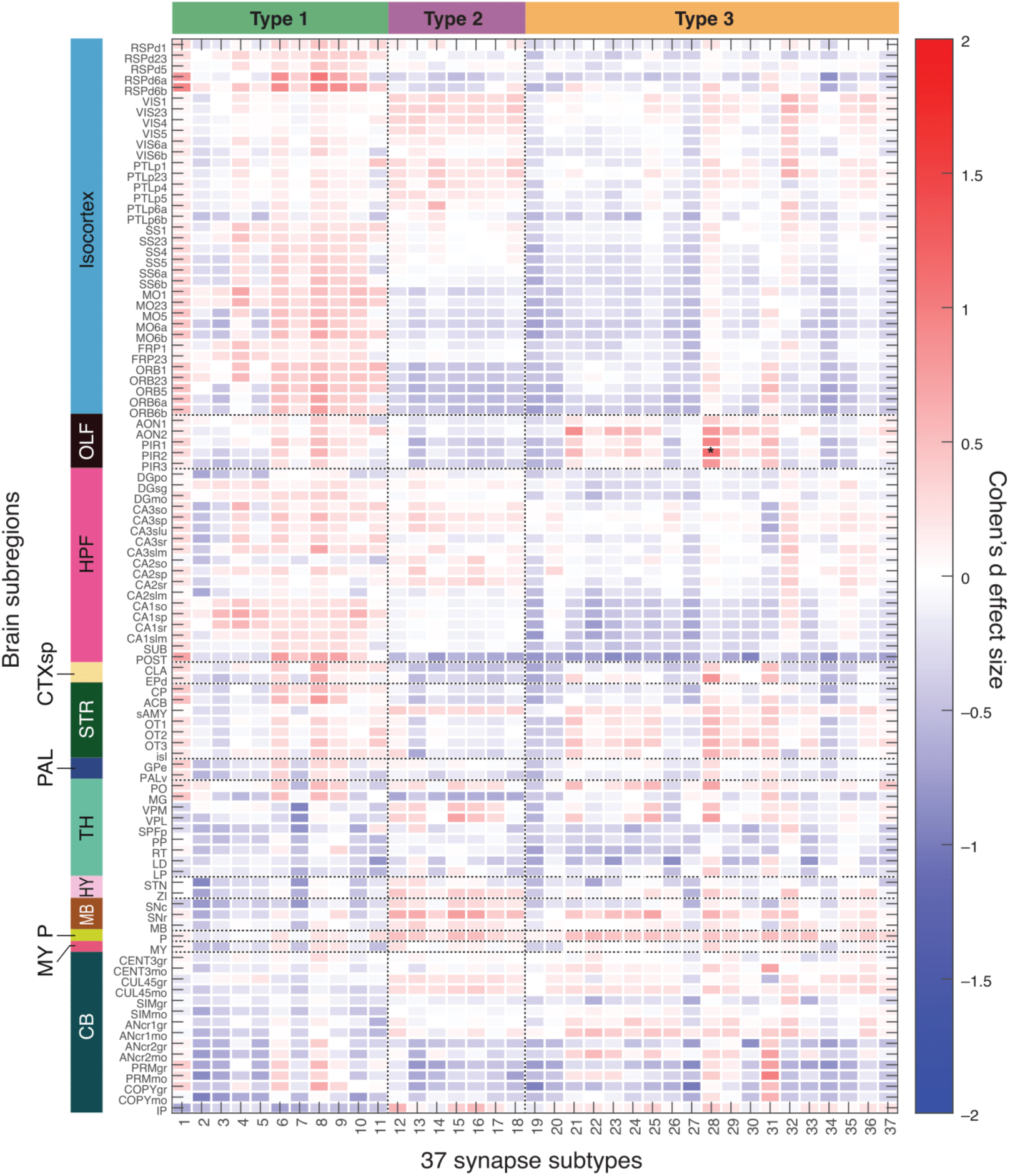
Cohen’s *d* effect size values comparing synapse subtype densities between 3-month SC female and male mice across all brain subregions. Red indicates higher density in females. Asterisk denotes statistical significance (*p < 0.05); no significance was retained after Benjamini–Hochberg correction applied to Bayesian posterior probabilities, indicating no consistent sex differences in synaptome architecture under SC. For brain region abbreviations see supplementary material ‘Brain region list’.

**Fig. S4).**
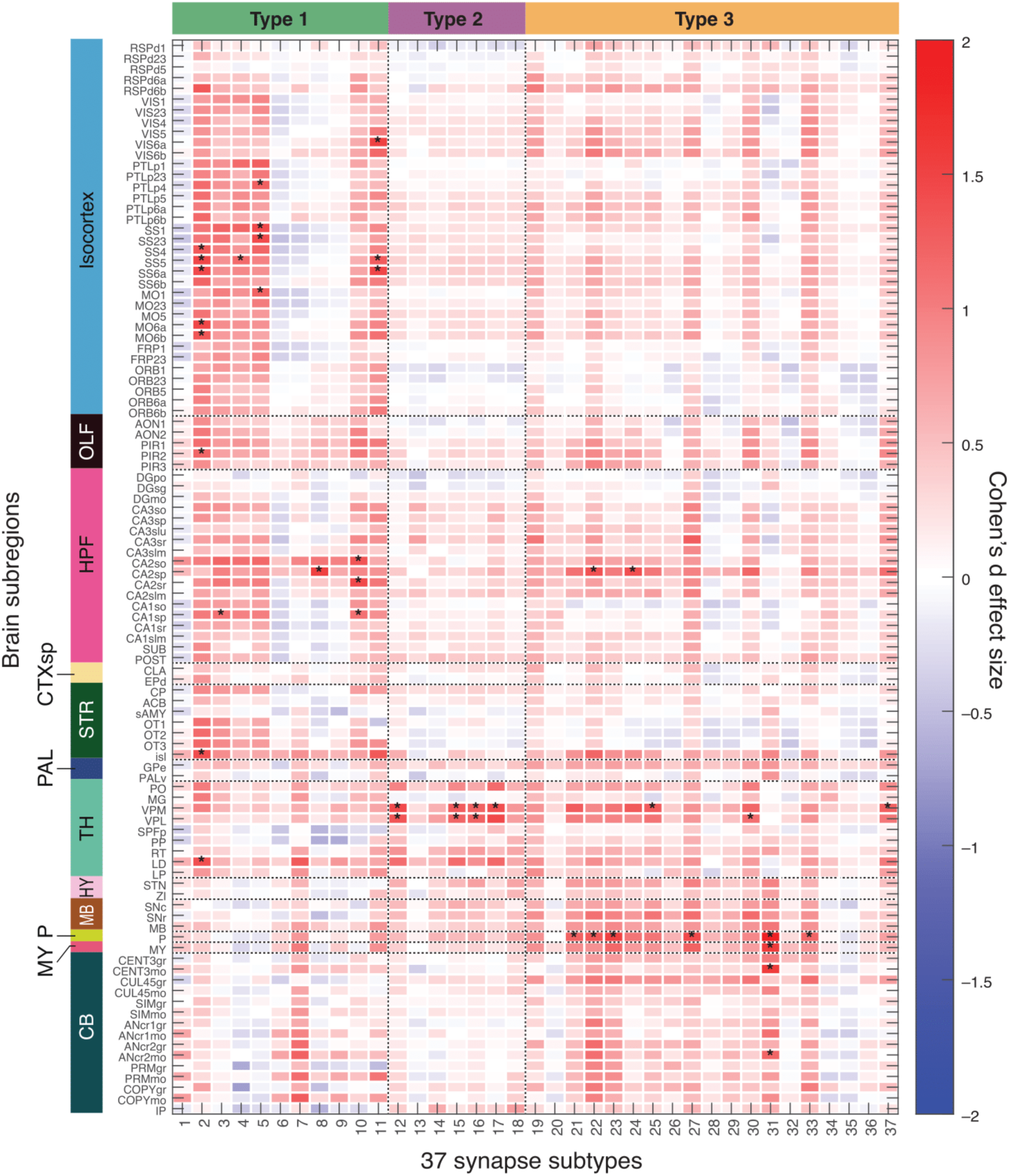
Cohen’s *d* effect size values comparing synapse subtype densities between developmental EE female and male mice. Red indicates higher density in females. Asterisk denotes statistical significance (*p < 0.05); no significance was retained after Benjamini–Hochberg correction applied to Bayesian posterior probabilities, indicating no large sex differences in synaptome architecture under EE. However, trends indicate that developmental EE may trigger some sex-dependent changes in synaptome architecture. For brain region abbreviations see supplementary material ‘Brain region list’.

**Fig. S5).**
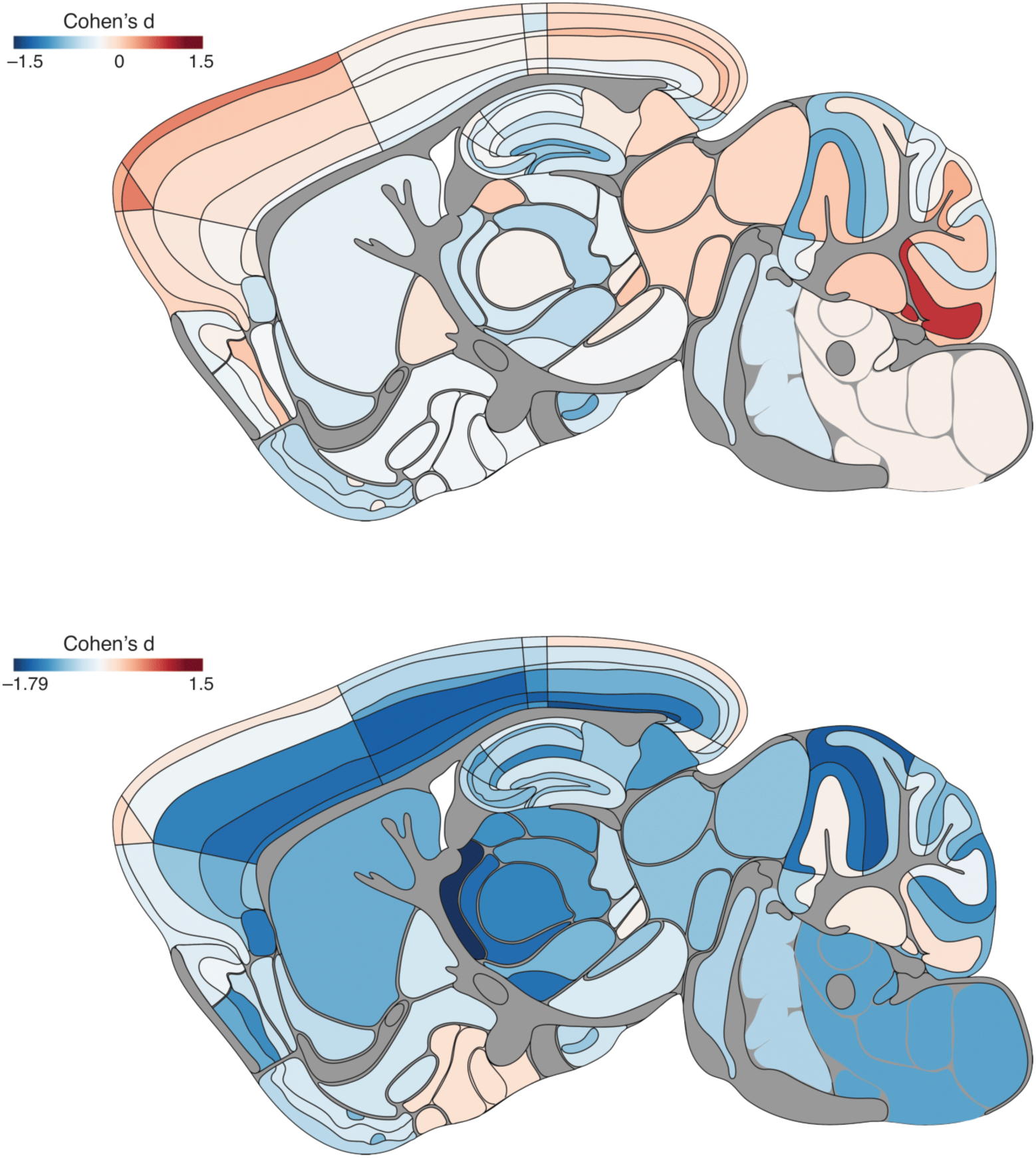
Parasagittal brain maps comparing total synapse density with Cohen’s *d* effect size between developmental EE and SC mice, split by sex. Top: EE vs. SC females; bottom: EE vs. SC males. Red indicates higher density in EE mice. See Fig. S29 for region key.

**Fig. S6).**
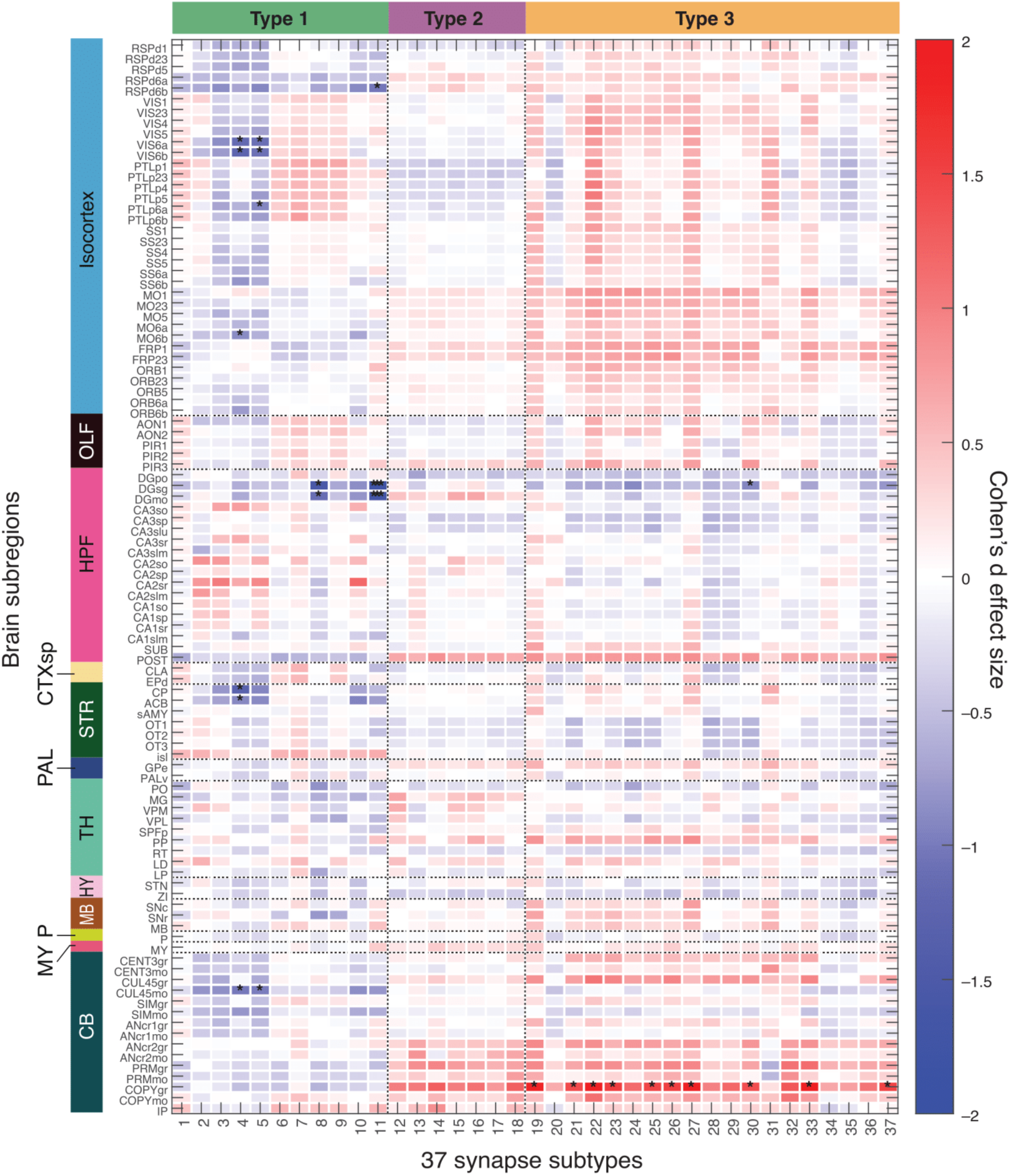
Cohen’s d effect size values comparing synapse subtype densities between developmental EE and SC females. Red indicates higher density in EE mice. Asterisks denote statistical significance (*p < 0.05; ***p < critical p after Benjamini–Hochberg correction applied to Bayesian posterior probabilities). Only subtype 11 was found to be significant after correction (p < 0.00047). EE females showed non-significant trends of higher Type 3 subtypes and lower Type 1 subtypes in the isocortex and cerebellum. For brain region abbreviations see supplementary material ‘Brain region list’.

**Fig. S7).**
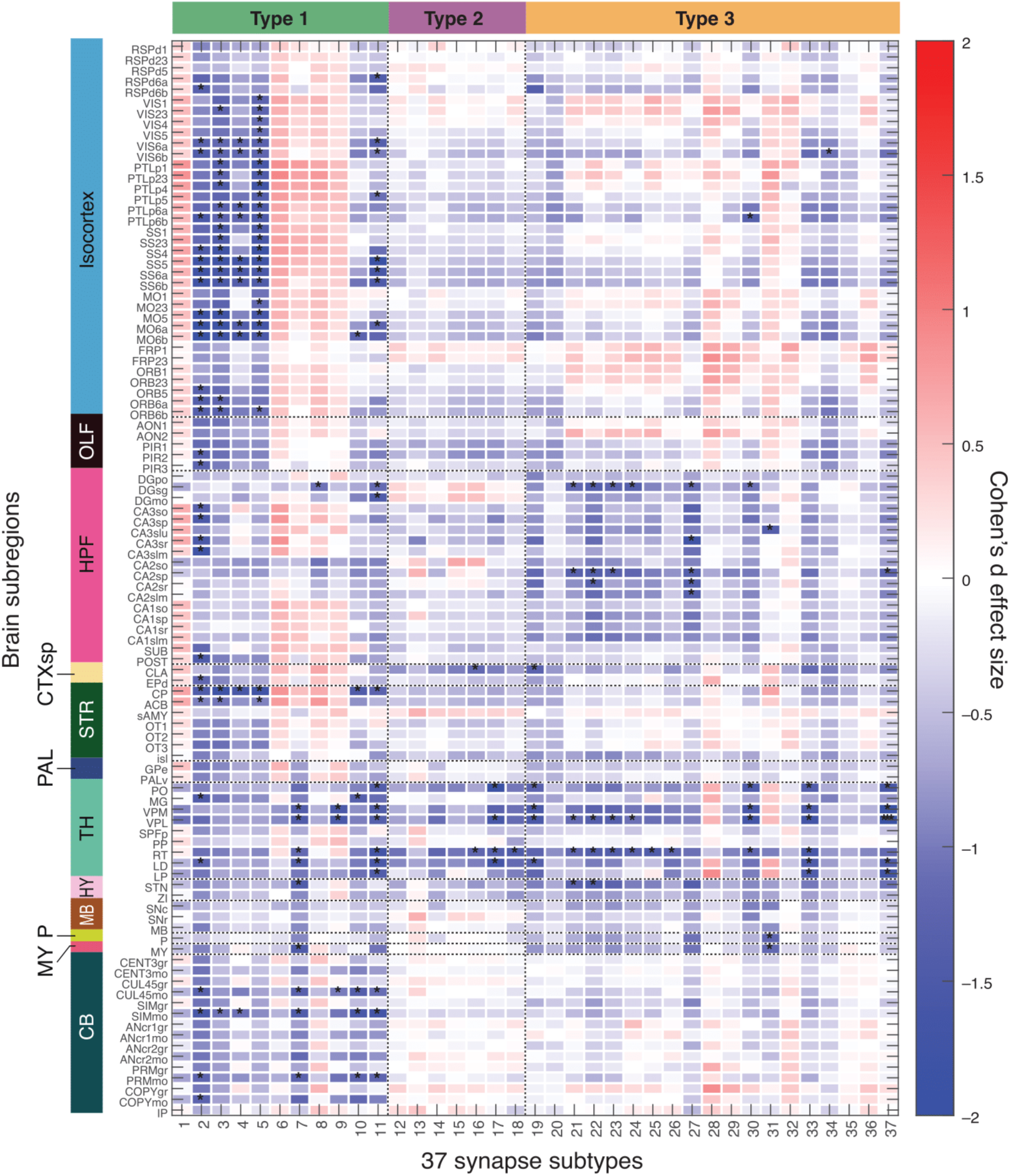
Cohen’s *d* effect size values comparing synapse subtype densities between developmental EE and SC males. Red indicates higher density in EE mice. Asterisks denote statistical significance (*p < 0.05; ***p < critical p after Benjamini–Hochberg correction applied to Bayesian posterior probabilities). Subtype 37 significance remained after correction (p < 0.0009). EE males exhibited widespread reduction in a subset of Type 1 subtypes, with additional decreases in Type 3 subtypes in hippocampus and thalamus. For brain region abbreviations see supplementary material ‘Brain region list’.

**Fig. S8).**
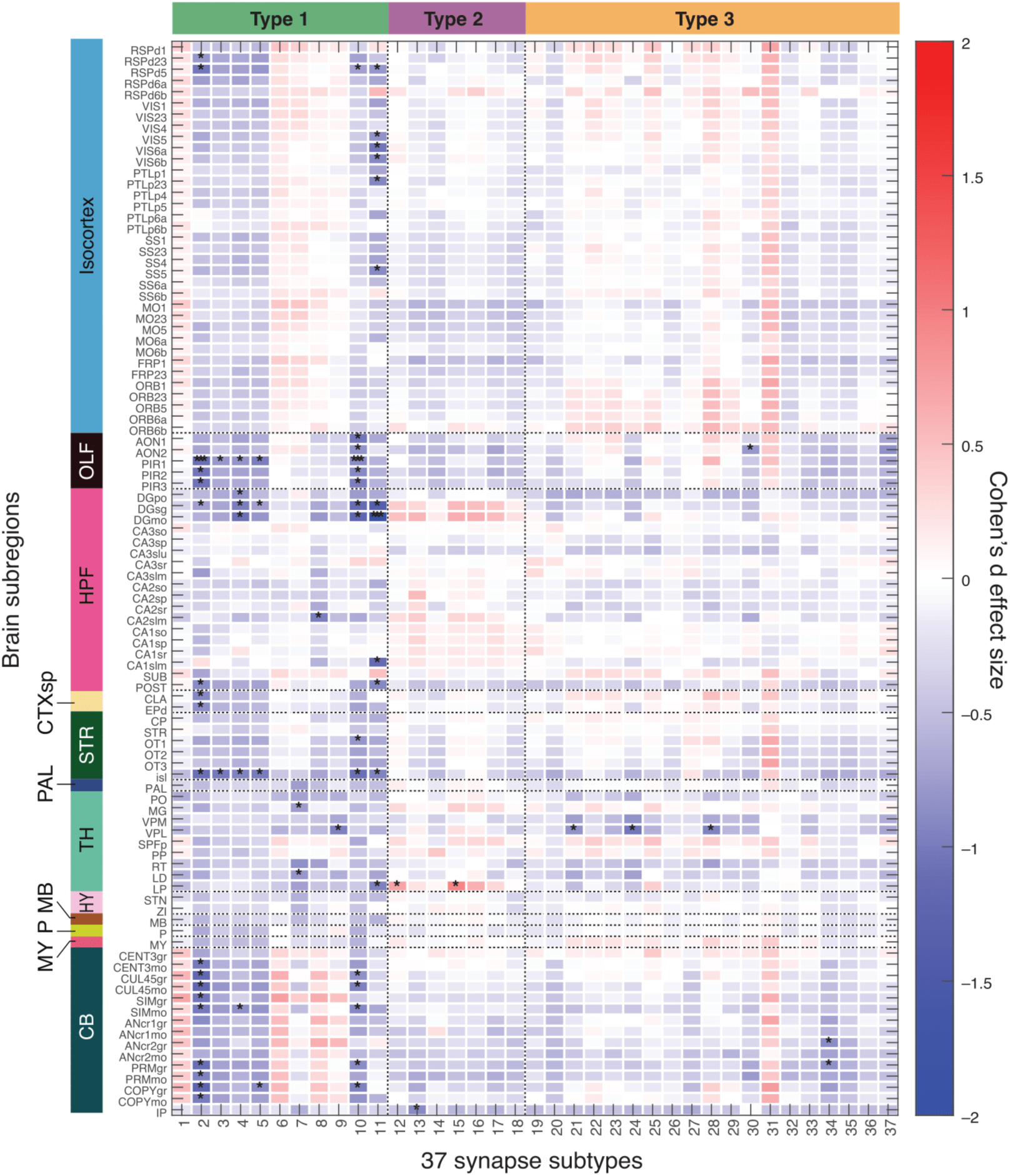
Cohen’s *d* effect size values comparing synapse subtype densities between adult EE and SC mice. Red indicates higher density in EE mice. Asterisks denote statistical significance (*p < 0.05; ***p < critical p after Benjamini–Hochberg correction applied to Bayesian posterior probabilities). Subtypes 2, 10, and 11 showed significance after correction (p < 0.0004). For brain region abbreviations see supplementary material ‘Brain region list’.

**Fig. S9).**
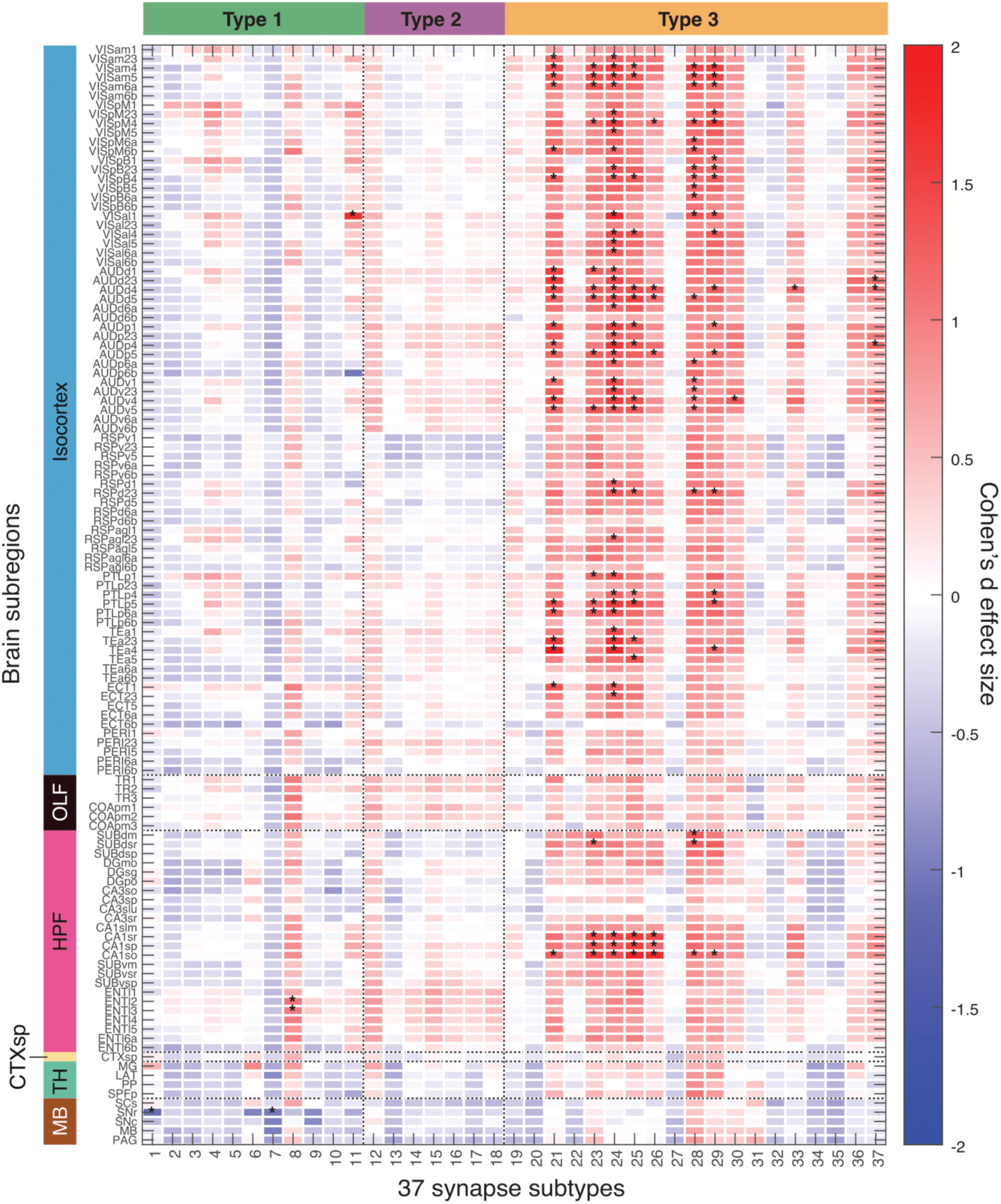
Cohen’s *d* effect size values comparing synapse subtype densities across the contralateral hemisphere between MD and control mice. Red indicates higher density in MD mice. Asterisk denotes statistical significance (*p < 0.05); no significance was retained after Benjamini–Hochberg correction applied to Bayesian posterior probabilities. For brain region abbreviations see supplementary material ‘Brain region list’.

**Fig. S10).**
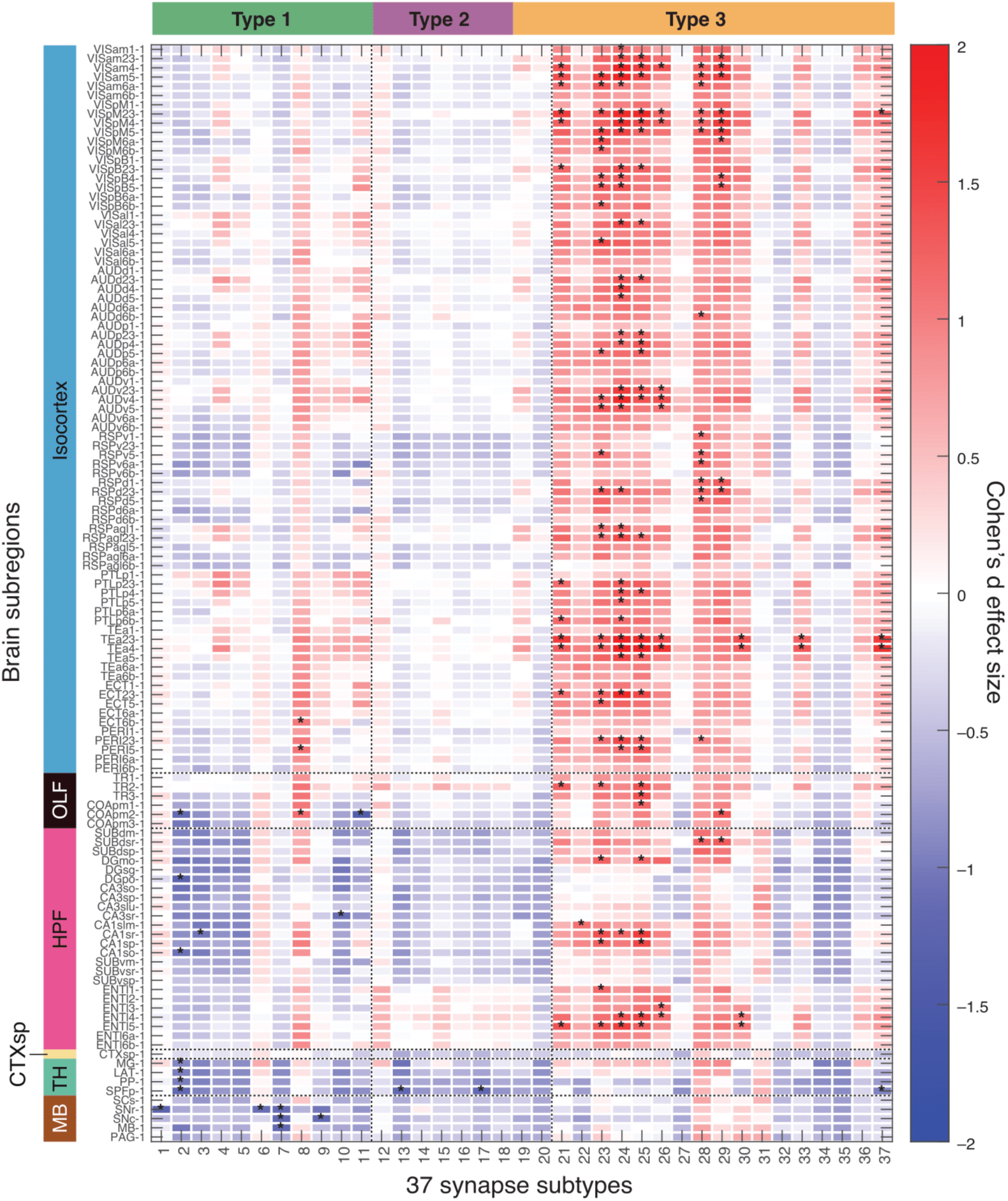
Cohen’s *d* effect size values comparing synapse subtype densities across the ipsilateral hemisphere between MD and control mice. Red indicates higher density in MD mice. Asterisk denotes statistical significance (*p < 0.05); no significance was retained after Benjamini–Hochberg correction applied to Bayesian posterior probabilities. A trend toward lower Type 1 and Type 2 subtype densities was observed in the hippocampus, thalamus, and midbrain—distinct from the contralateral hemisphere. For brain region abbreviations see supplementary material ‘Brain region list’.

**Fig. S11).**
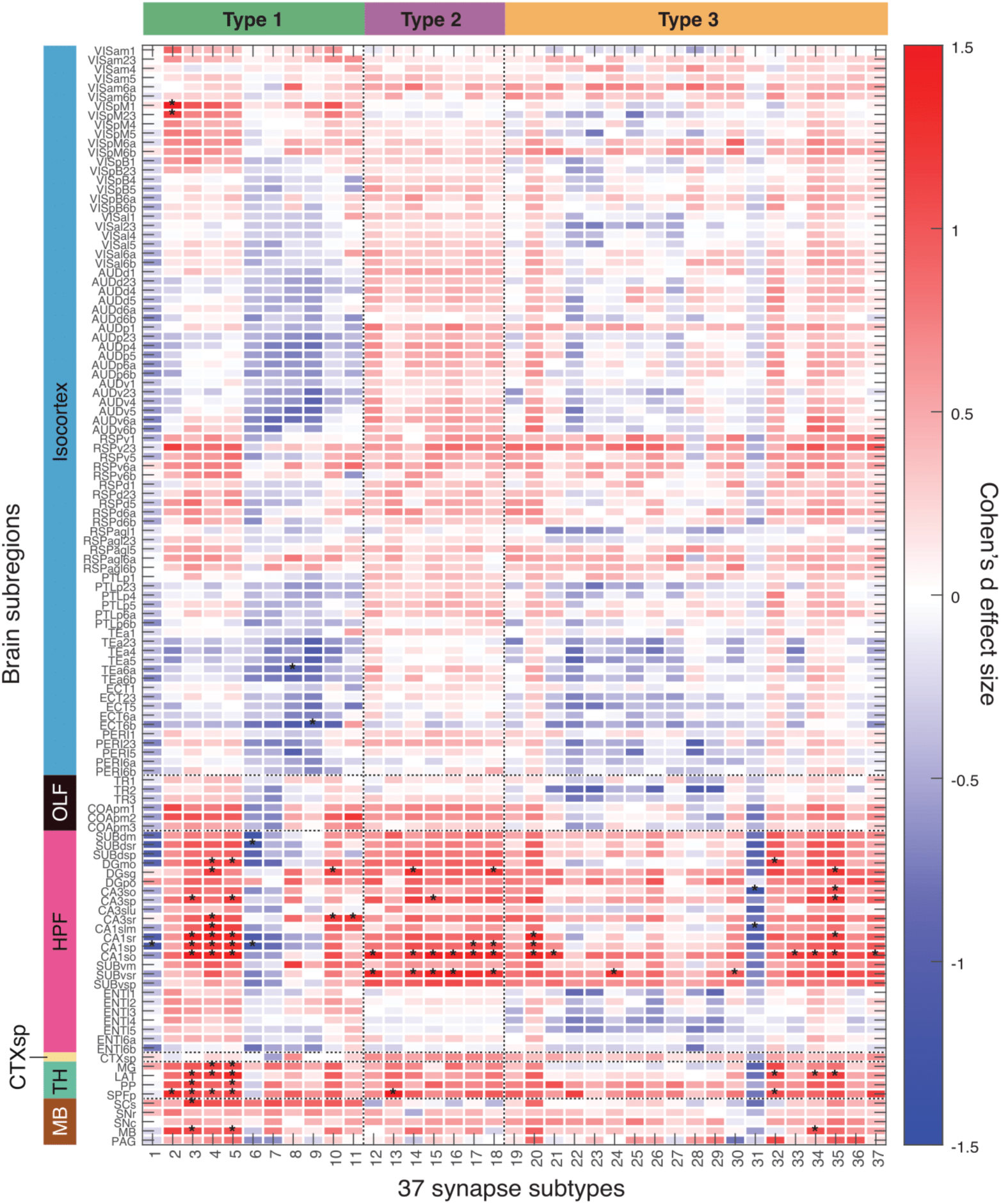
Cohen’s *d* effect size values comparing synapse subtype densities across the contralateral hemisphere, normalised by summed hemisphere values, between MD and control mice. Red indicates subtypes that were relatively more abundant in the contralateral hemisphere of MD mice than in controls. Asterisk denotes statistical significance (*p < 0.05); no significance was retained after Benjamini–Hochberg correction applied to Bayesian posterior probabilities. A subset of subtypes— unrestricted by protein class—were proportionally higher in the visual cortex, retrosplenial cortex, hippocampus, thalamus, and midbrain of the contralateral hemisphere of MD mice. For brain region abbreviations see supplementary material ‘Brain region list’.

**Fig. S12).**
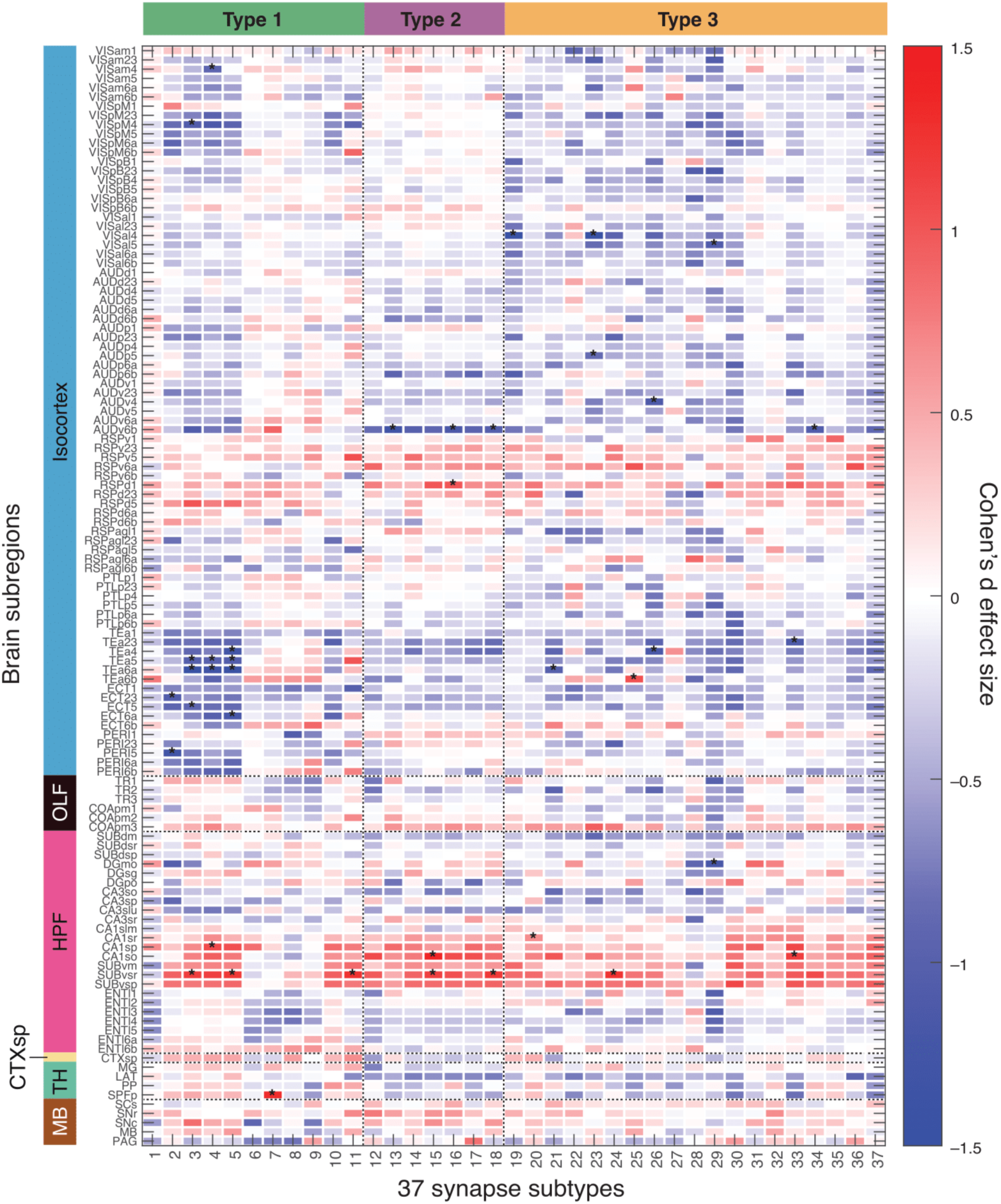
Cohen’s *d* effect size values comparing inter-hemispheric differences in subtype densities, normalised by summed hemisphere values, between MD and control mice. Red indicates greater asymmetry in MD mice than in controls. Asterisk denotes statistical significance (*p < 0.05); no significance was retained after Benjamini– Hochberg correction applied to Bayesian posterior probabilities. MD-induced shifts in the two hemispheres reduced asymmetry in many isocortical regions, while increasing asymmetry in the CA1 and subiculum. For brain region abbreviations see supplementary material ‘Brain region list’.

**Fig. S13).**
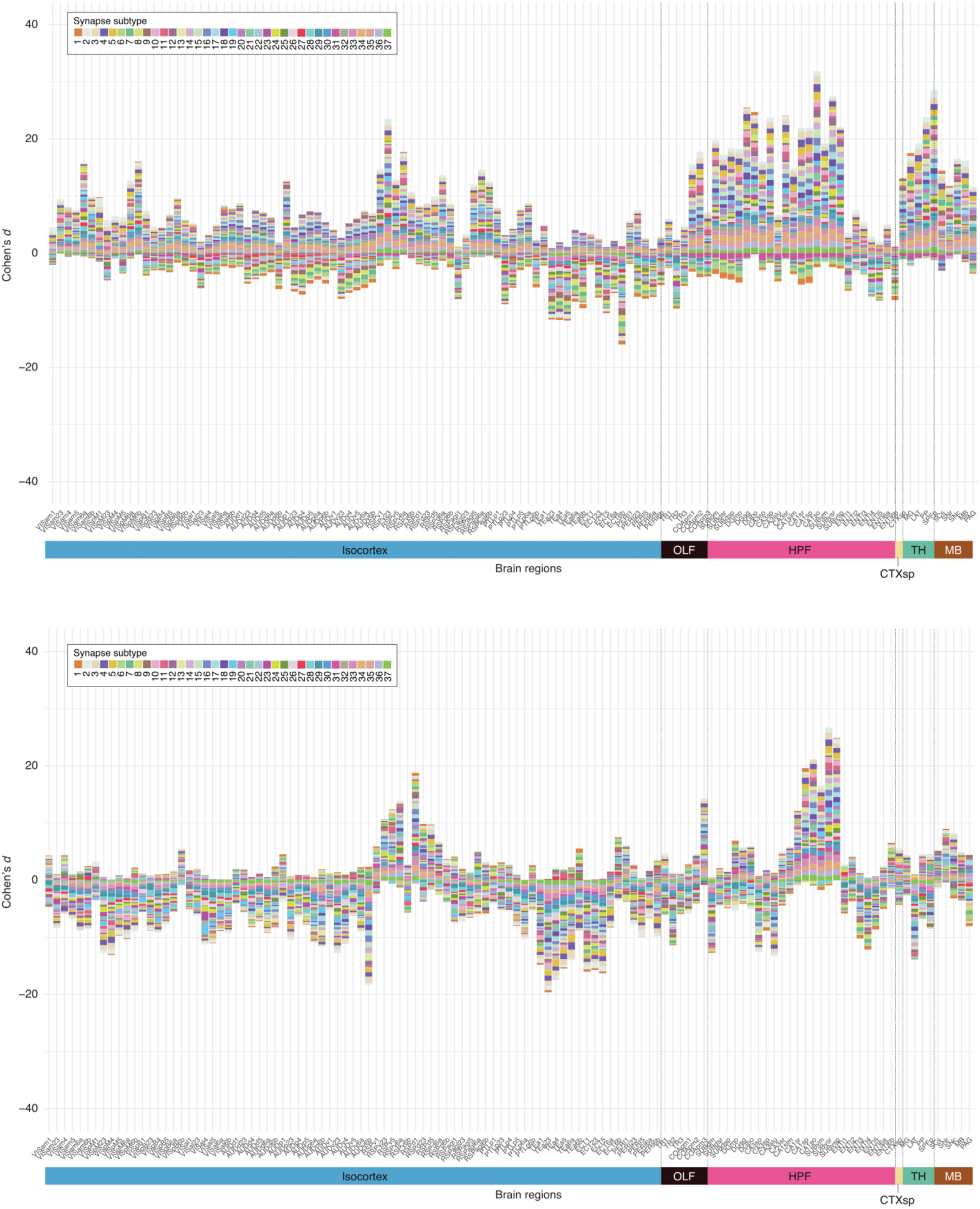
Top: Stacked bar graph of Cohen’s d values for 37 synapse subtypes in the contralateral (deprived) hemisphere, comparing MD and control mice. Subregion values were normalized by dividing contralateral densities by the sum of ipsilateral and contralateral densities. Positive values indicate increased relative contralateral density in MD. Bottom: Stacked bar graph of Cohen’s d values for 37 synapse subtypes comparing interhemispheric differences between MD and control mice. The interhemispheric difference for each subtype and subregion was calculated by dividing the absolute difference between contralateral and ipsilateral densities by the sum of ipsilateral and contralateral densities. Positive values indicate increased interhemispheric difference with MD. For brain region abbreviations see supplementary material ‘Brain region list’.

**Fig. S14).**
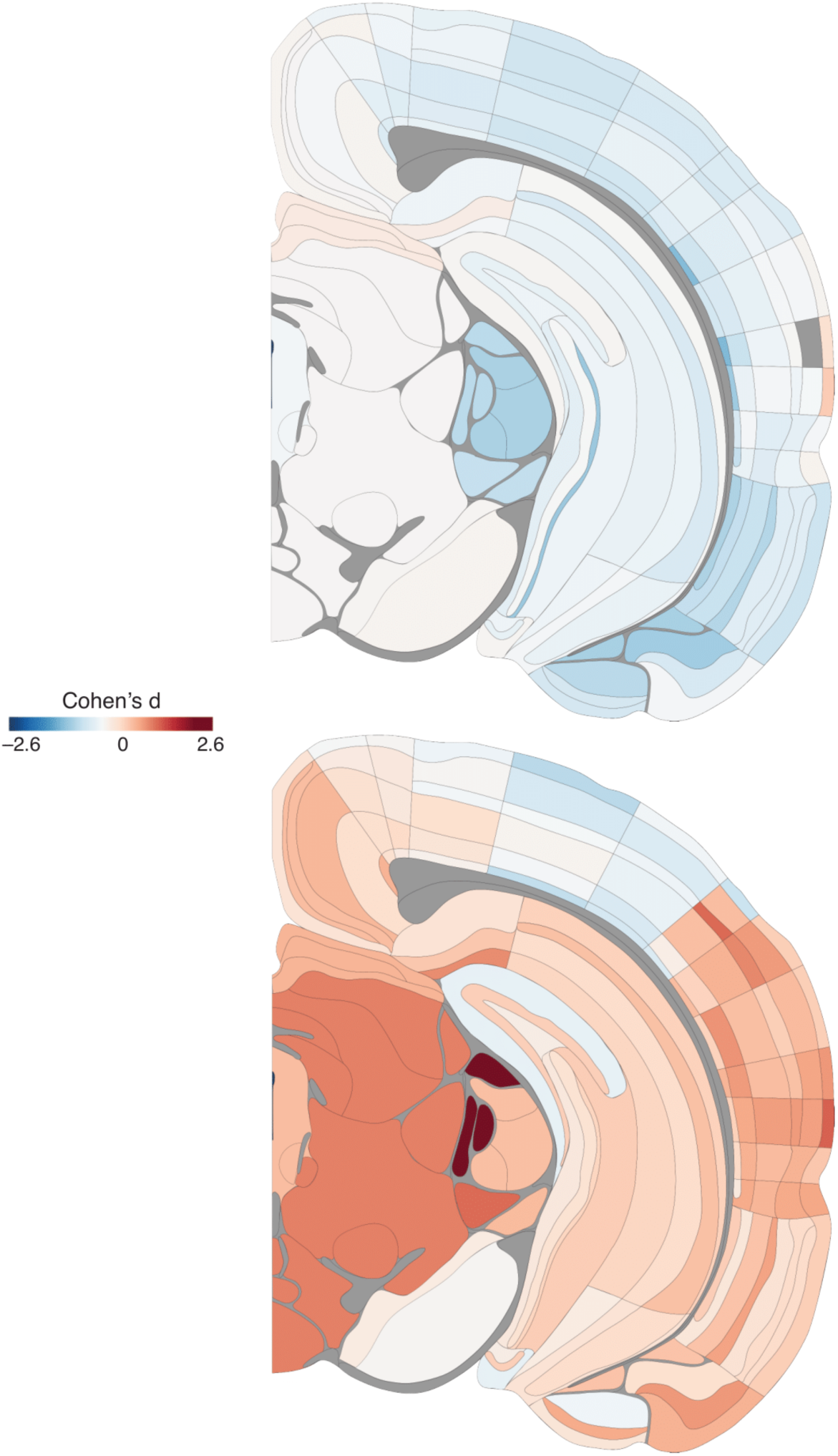
Coronal brain maps comparing total synapse density in the contralateral hemisphere with Cohen’s *d* effect size between control females and males (top), and MD females and males (bottom). Red indicates higher density in females. See Fig. S28 for region key.

**Fig. S15).**
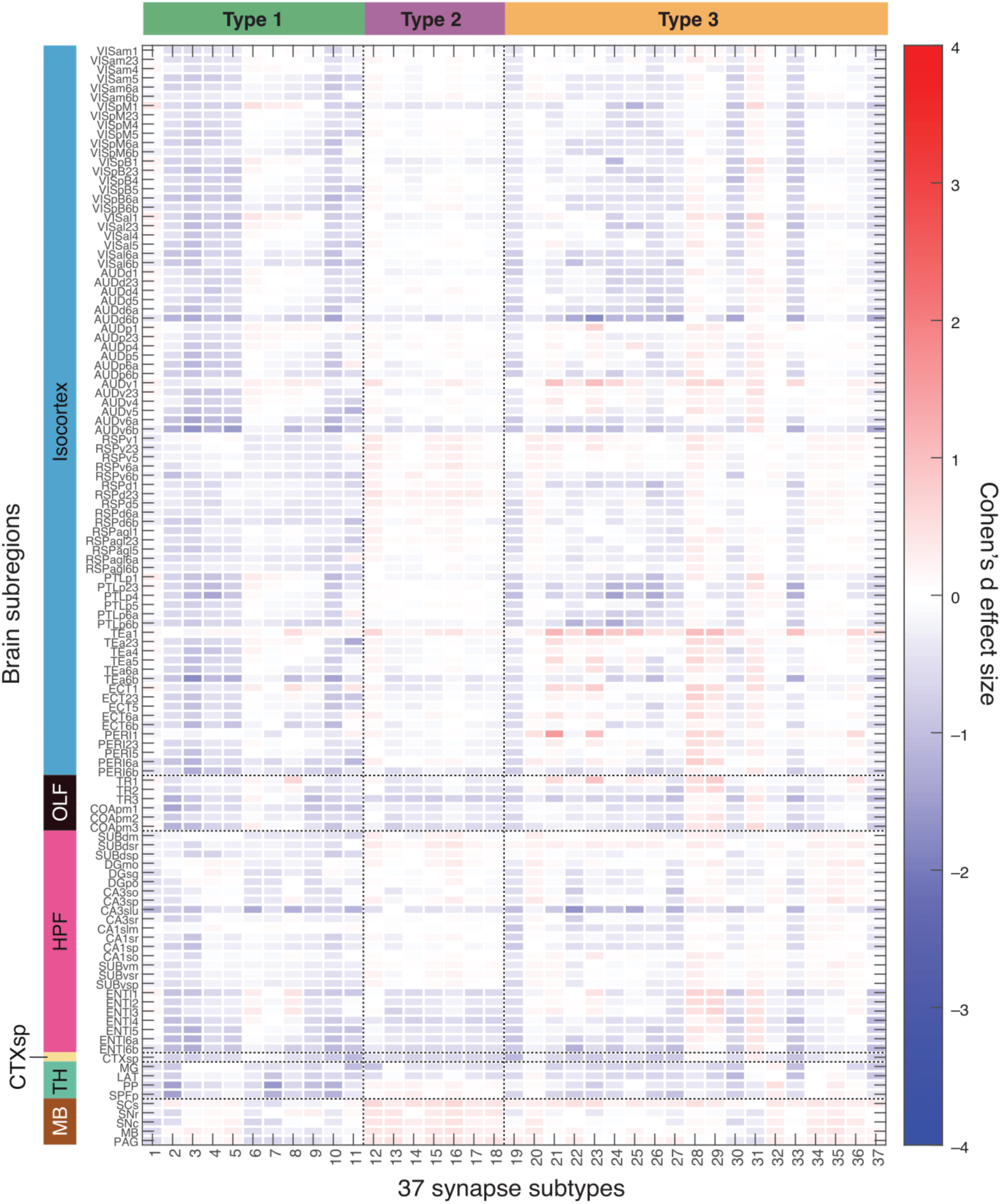
Cohen’s *d* effect size values comparing synapse subtype densities across the contralateral hemisphere between control females and males. Red indicates higher density in females. No significance before or after correction. A trend of lower Type 1 subtypes in the isocortex of female controls was observed. For brain region abbreviations see supplementary material ‘Brain region list’.

**Fig. S16).**
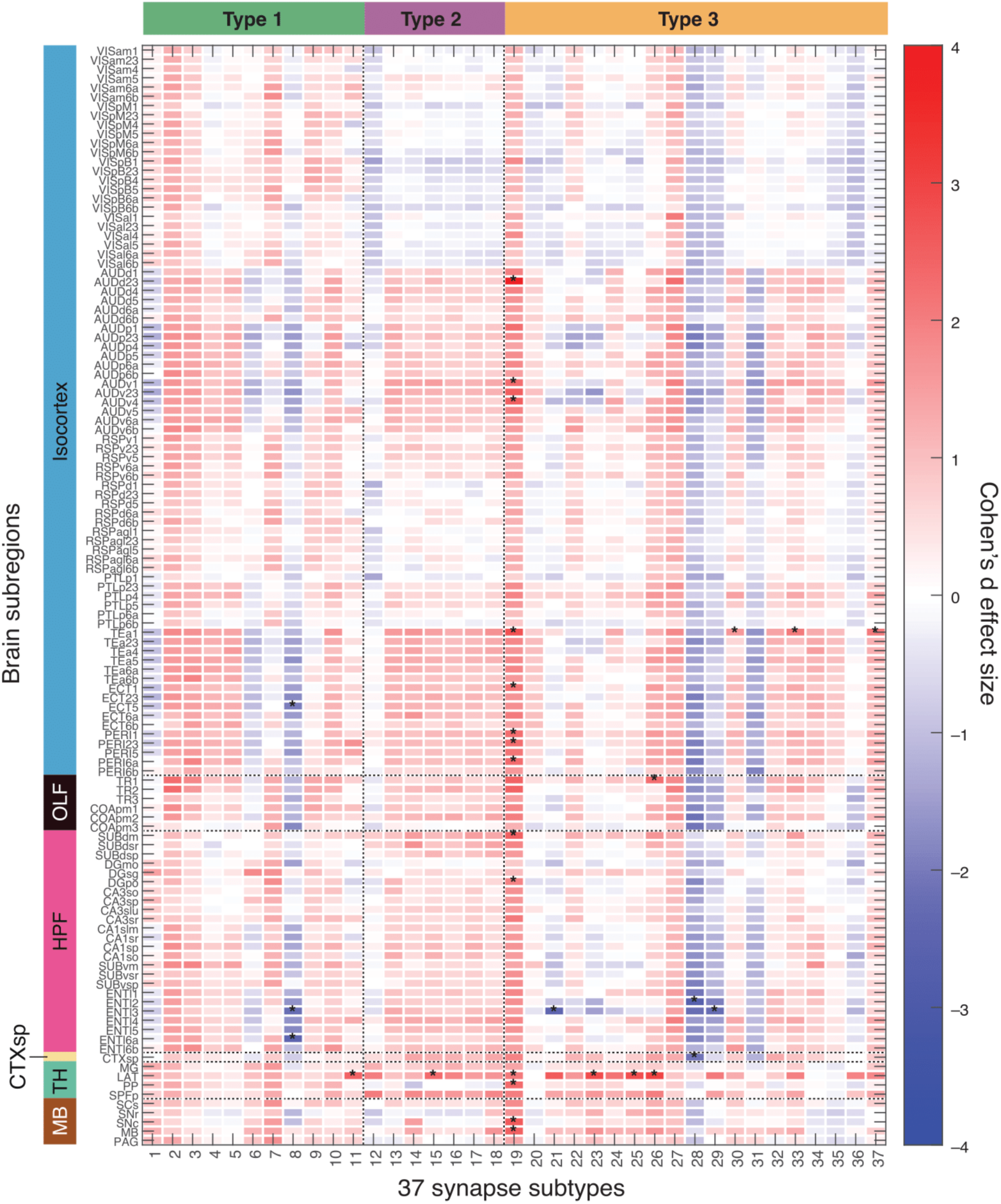
Cohen’s *d* effect size values comparing synapse subtype densities across the contralateral hemisphere between MD females and males. Red indicates higher density in females. Asterisk denotes statistical significance (*p < 0.05); no significance was retained after Benjamini–Hochberg correction applied to Bayesian posterior probabilities. Subtypes higher in females were not restricted to specific synapse types. As in EE, MD appears to introduce sex-dependent synaptome shifts. For brain region abbreviations see supplementary material ‘Brain region list’.

**Fig. S17).**
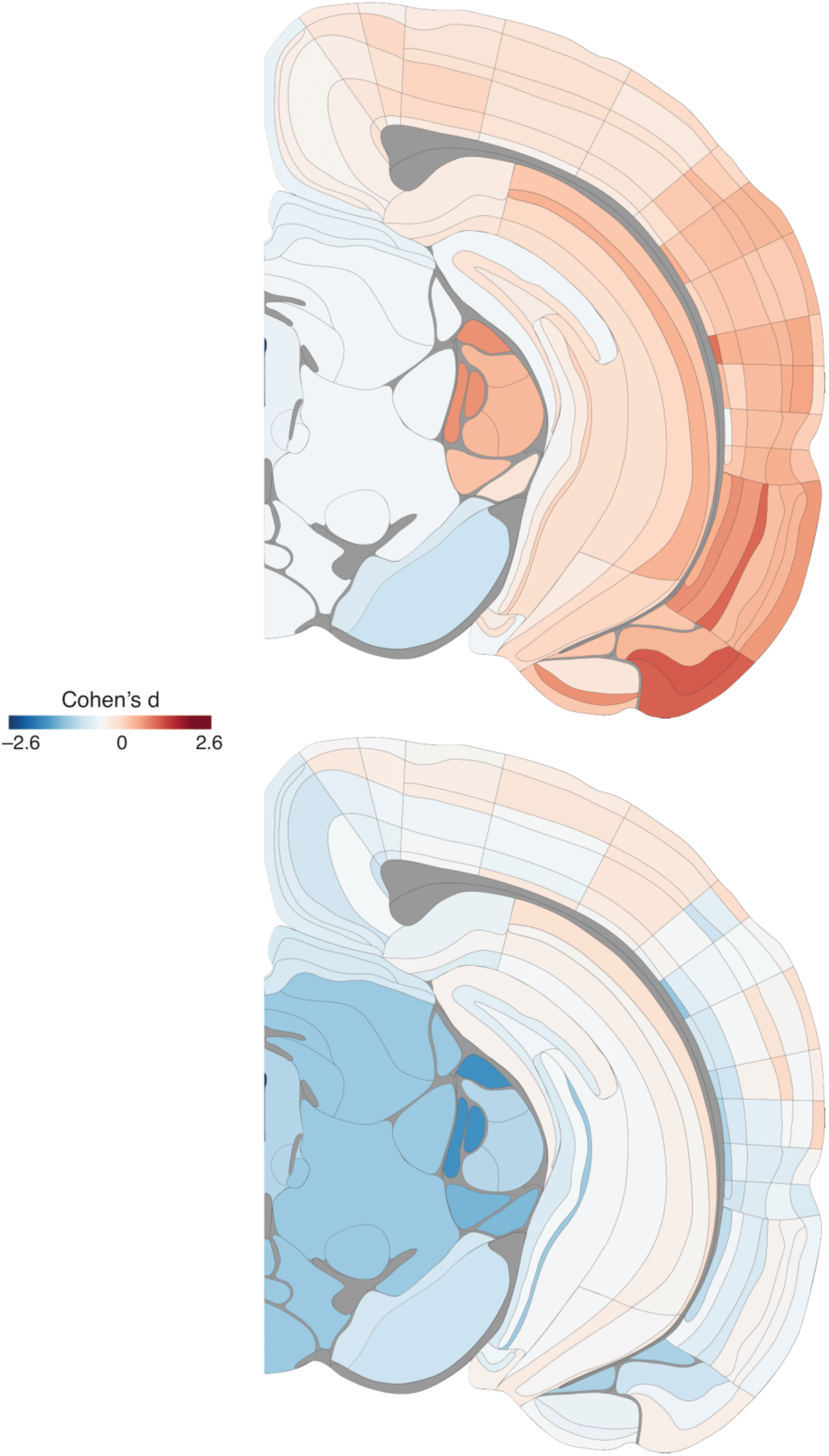
Coronal brain maps comparing total synapse density with Cohen’s *d* effect size between MD and control mice, split by sex. Top: MD vs. control females. Bottom: MD vs. control males. Red indicates higher density in MD mice. See Fig. S28 for region key.

**Fig. S18).**
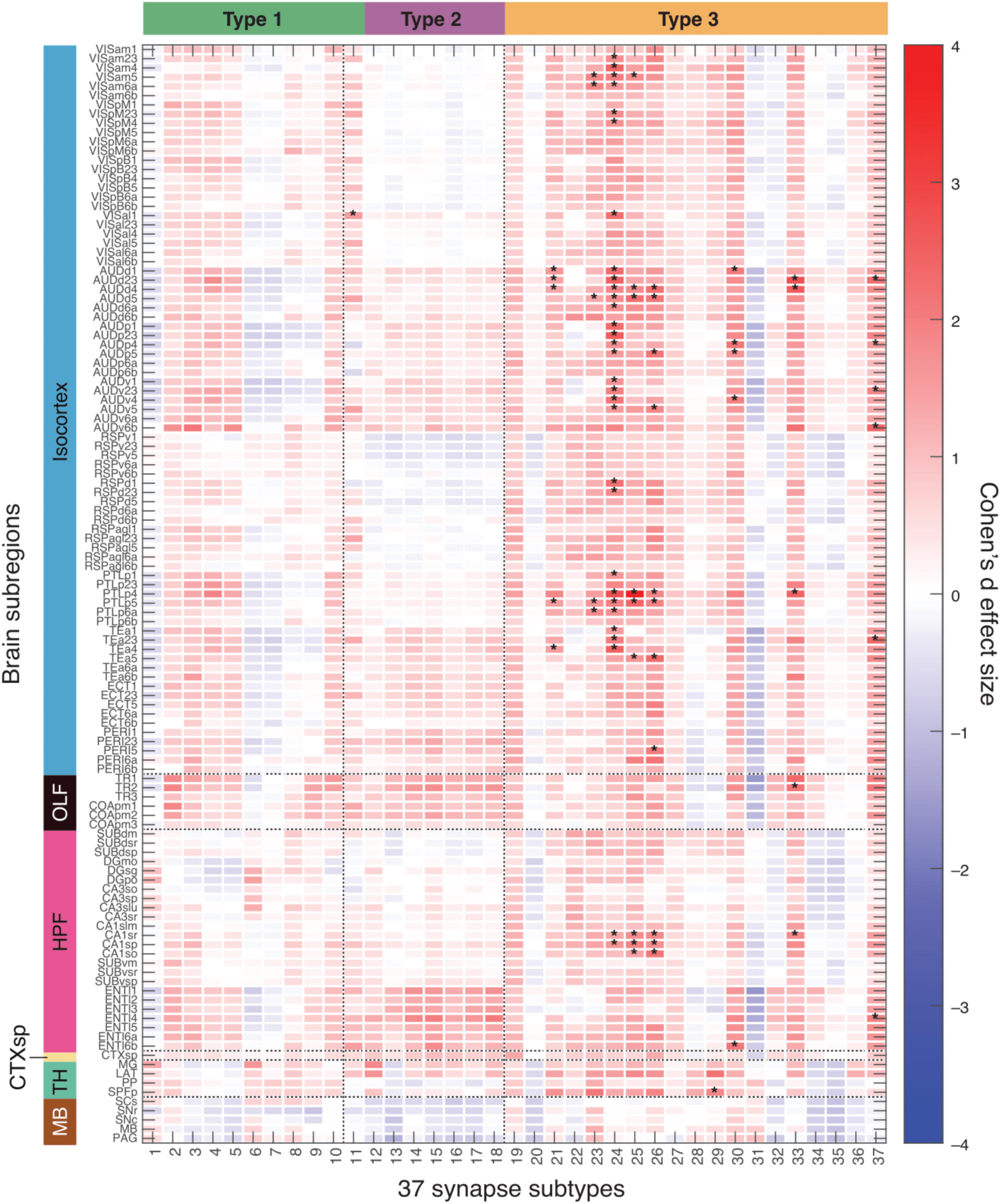
Cohen’s *d* effect size values comparing synapse subtype densities across the contralateral hemisphere between MD and control females. Red indicates higher density in MD mice. Asterisk denotes statistical significance (*p < 0.05); no significance was retained after Benjamini–Hochberg correction applied to Bayesian posterior probabilities. A subset of subtypes showed elevated density in MD females, independent of synapse type. For brain region abbreviations see supplementary material ‘Brain region list’.

**Fig. S19).**
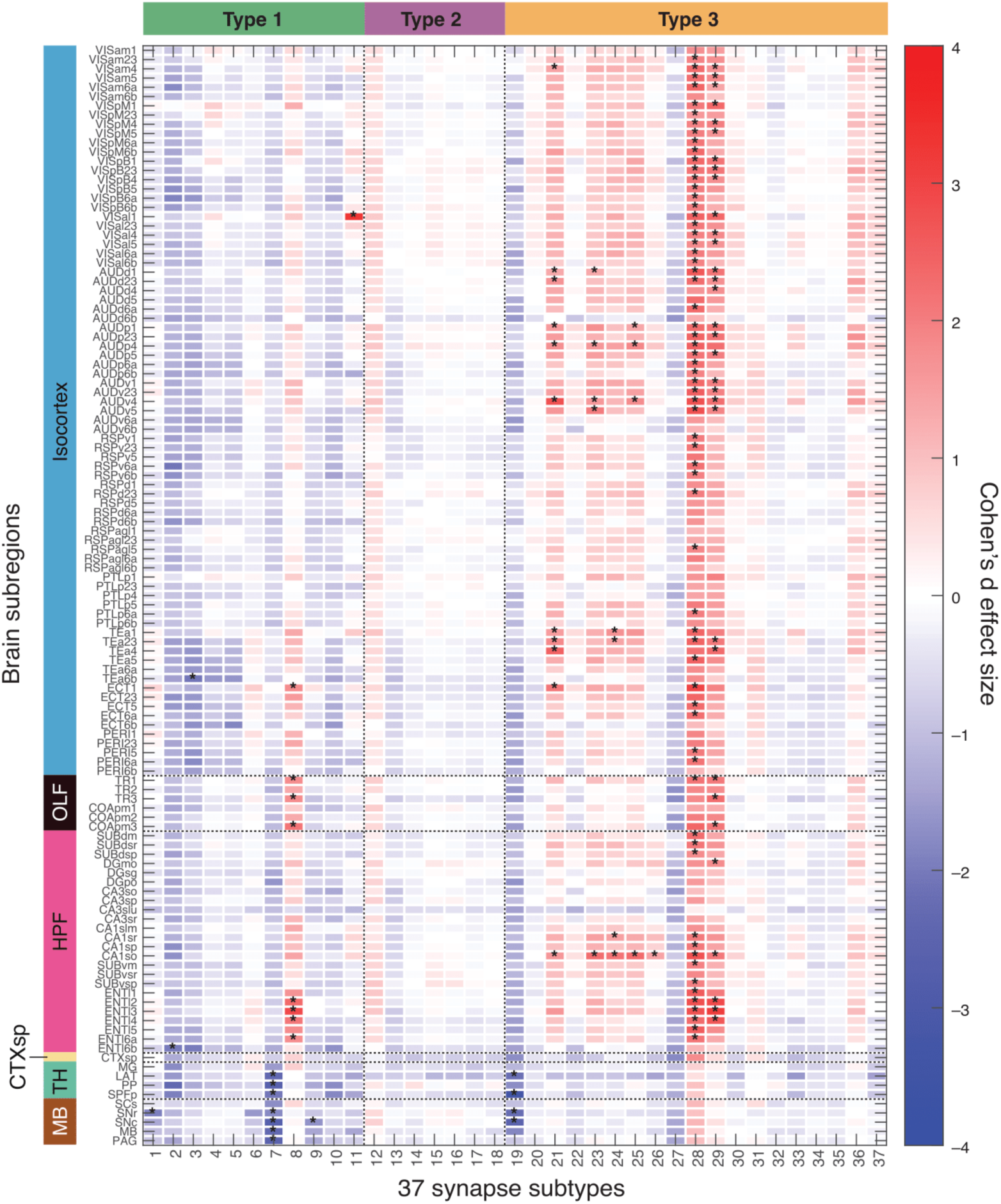
Cohen’s *d* effect size values comparing synapse subtype densities across the contralateral hemisphere between MD and control males. Red indicates higher density in MD mice. Asterisk denotes statistical significance (*p < 0.05); no significance was retained after Benjamini–Hochberg correction applied to Bayesian posterior probabilities. Subtypes 28 and 29 showed higher density in the cortex of MD males. MD males showed reduced density of several Type 1 subtypes across the brain, alongside significant increases in select Type 3 subtypes. For brain region abbreviations see supplementary material ‘Brain region list’.

**Fig. S20).**
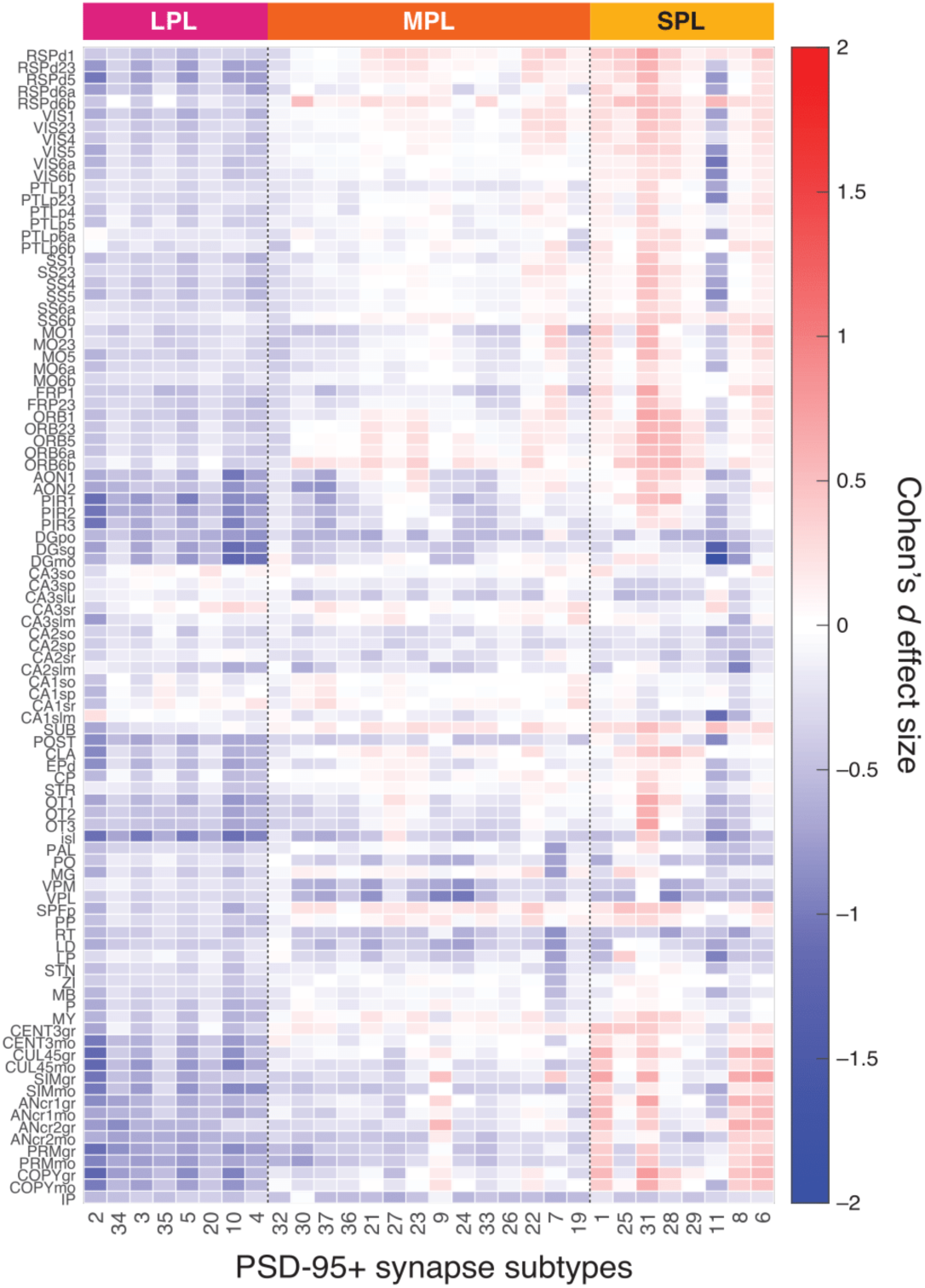
Cohen’s *d* effect size heatmaps showing EE-driven shifts in synapse subtypes categorised as long-protein lifetime (LPL) with adult EE paradigm. Similar to developmental EE, adult EE is associated with a preferential reduction in LPL synapses. For brain region abbreviations see supplementary material ‘Brain region list’.

**Fig. S21).**
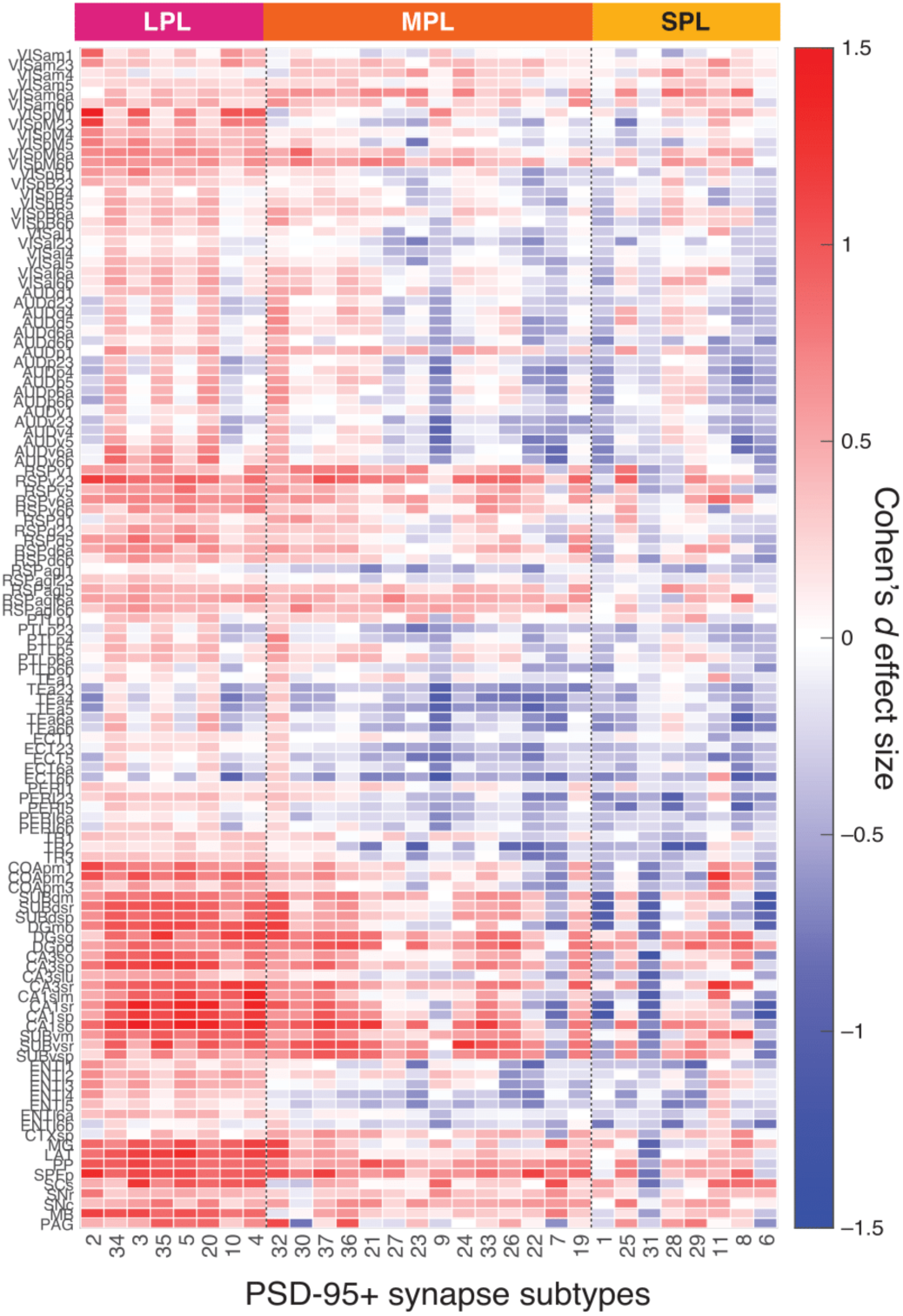
Cohen’s *d* effect size heatmaps showing hemisphere-specific, MD-driven shifts in synapse subtypes along a gradient of synapse protein lifetime. MD resulted in a higher density of long protein lifetime (LPL) synapses in the monocular visual area, retrosplenial areas, the hippocampus and the thalamus in the contralateral hemisphere, when normalised by summed hemisphere values. This lateralised reorganisation is reminiscent of changes observed following sleep deprivation (Koukaroudi et al., 2024). For brain region abbreviations see supplementary material ‘Brain region list’.

**Fig. S22).**
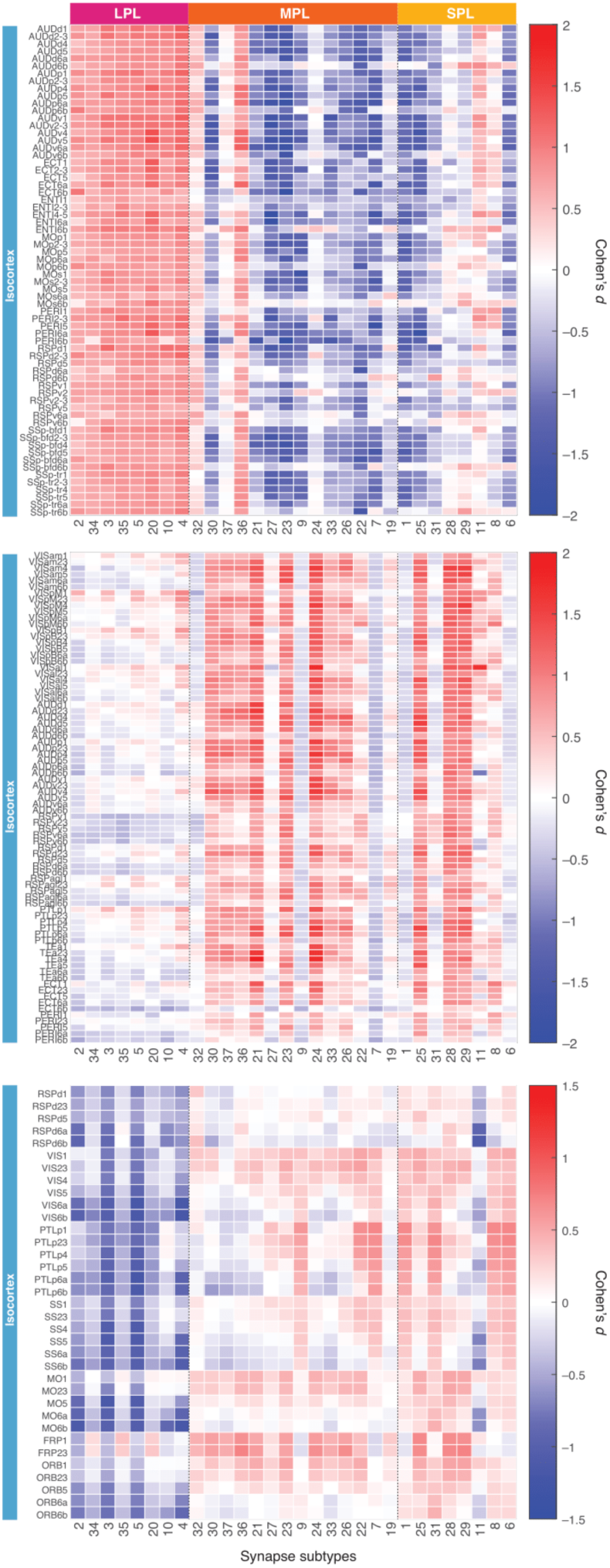
Regional effect sizes (Cohen’s d) for 30 PSD95⁺ synapse subtypes across the isocortex, ordered by protein lifetime, comparing sleep deprived mice and controls (top), MD mice and non-deprived controls (middle), and developmental EE and SC control mice (bottom). The sleep deprivation graph was reproduced from previously published data *(17)*. For brain region abbreviations see supplementary material ‘Brain region list’.

**Fig. S23).**
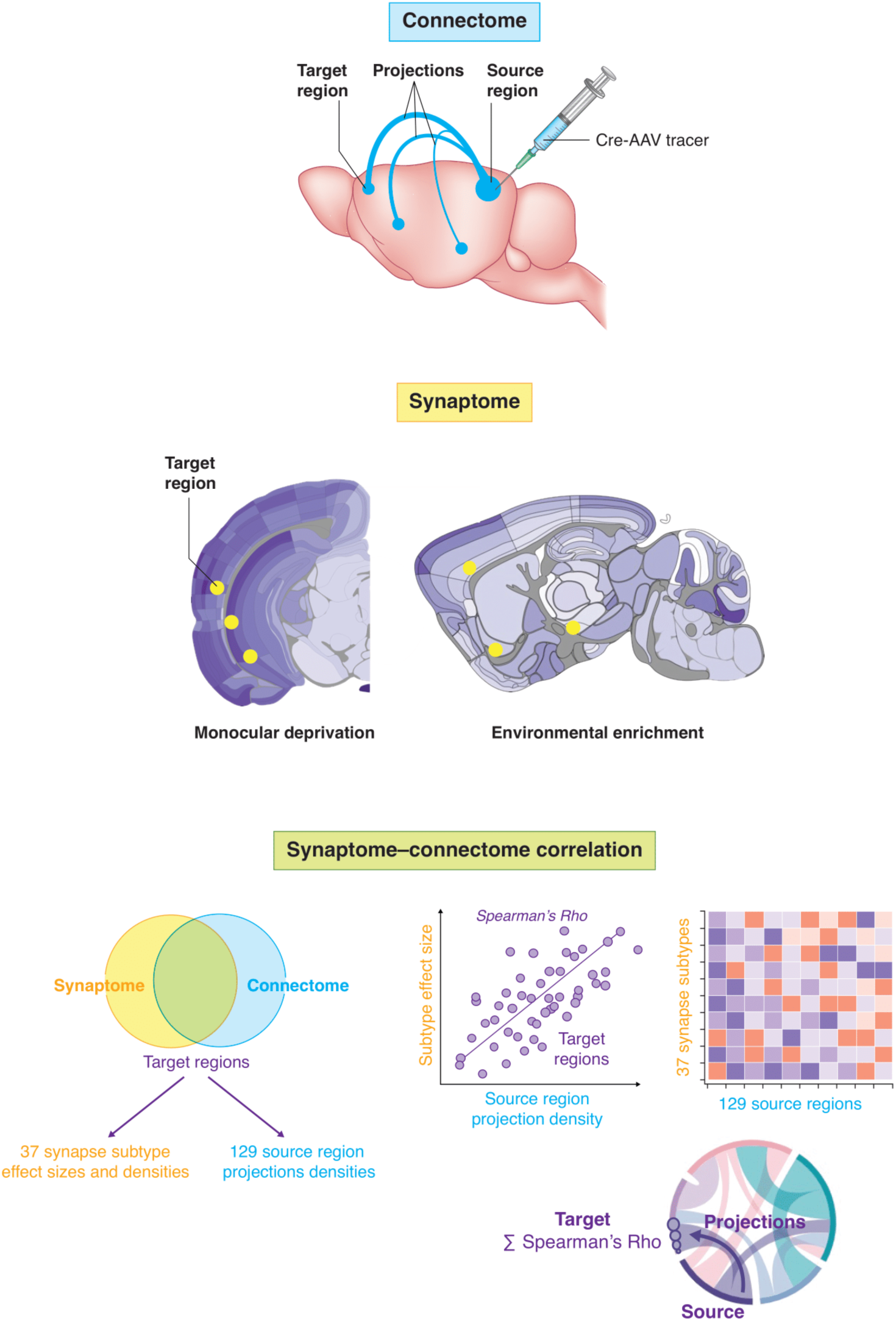
Overview of the synaptome–connectome correlation analysis. **Top:** Illustration of the tracing methodology used to produce the connectome datasets *(28)*, showing anterograde projection densities from each injected source region to all delineated target regions. **Middle:** Synaptome mapping provides regional synapse subtype density profiles and experimental effect sizes. **Bottom:** The two datasets are integrated across matched target regions to correlate projection strength from 129 source regions with either baseline subtype densities or Cohen’s d effect sizes for 37 subtypes. Representative outputs—scatter plot, heatmap, and network diagram—demonstrate result formats used in the main text.

**Fig. S24).**
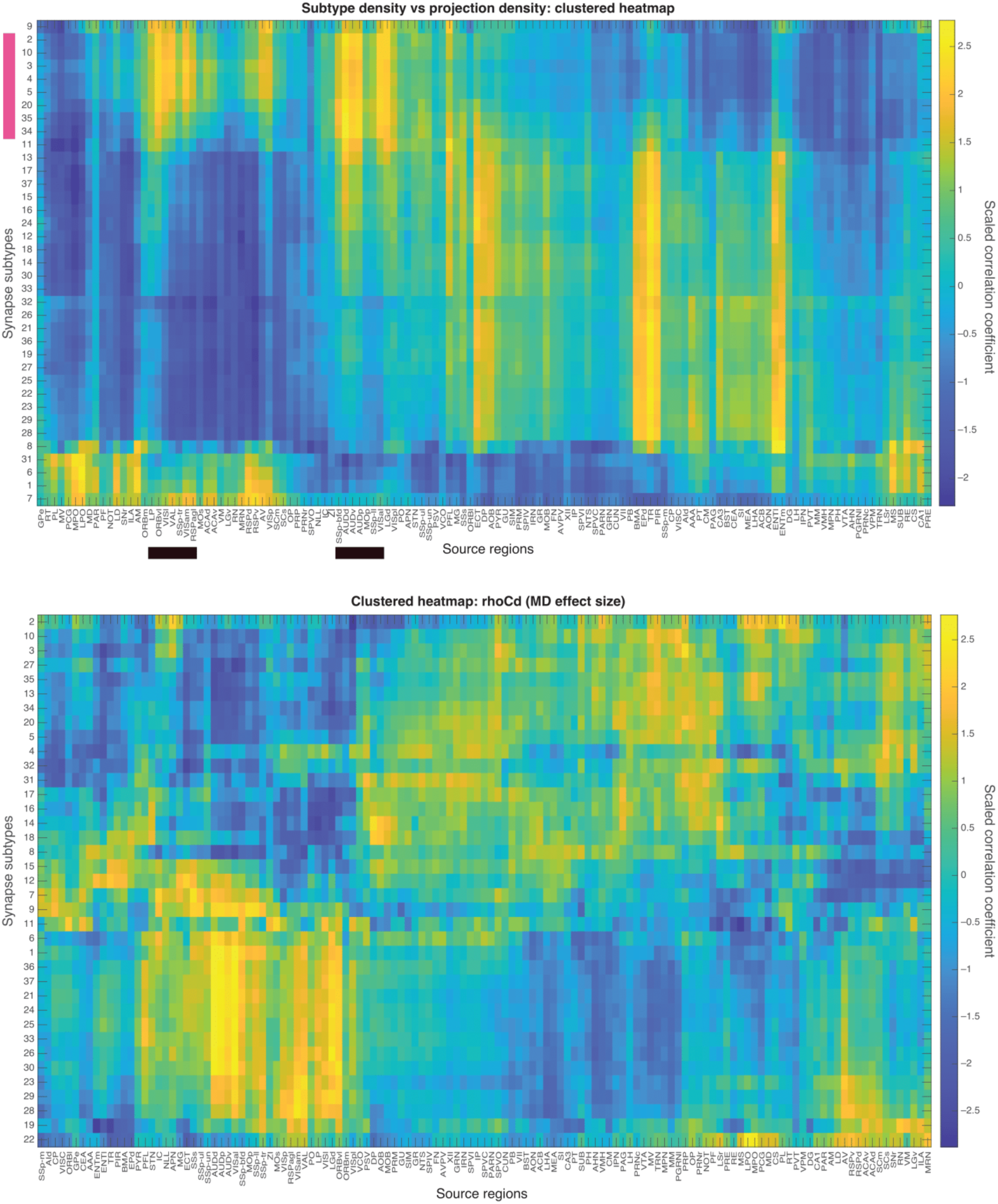
Top: Hierarchically clustered heatmaps of Spearman’s rho values correlating subtype densities (1-month control mice) with source region projection densities across target regions. The pink bar on the y-axis marks the cluster of long-protein lifetime synapses which positively correlate with several thalamocortical sensory areas, as indicated with the black bar on the x-axis. Bottom: Hierarchically clustered heatmaps of Spearman’s rho values correlating subtype effect size (MD paradigm) with source region projection densities across target regions. Rows represent 37 synapse subtypes; columns are anatomically defined source regions. For brain region abbreviations see supplementary material ‘Brain region list’.

**Fig. S25).**
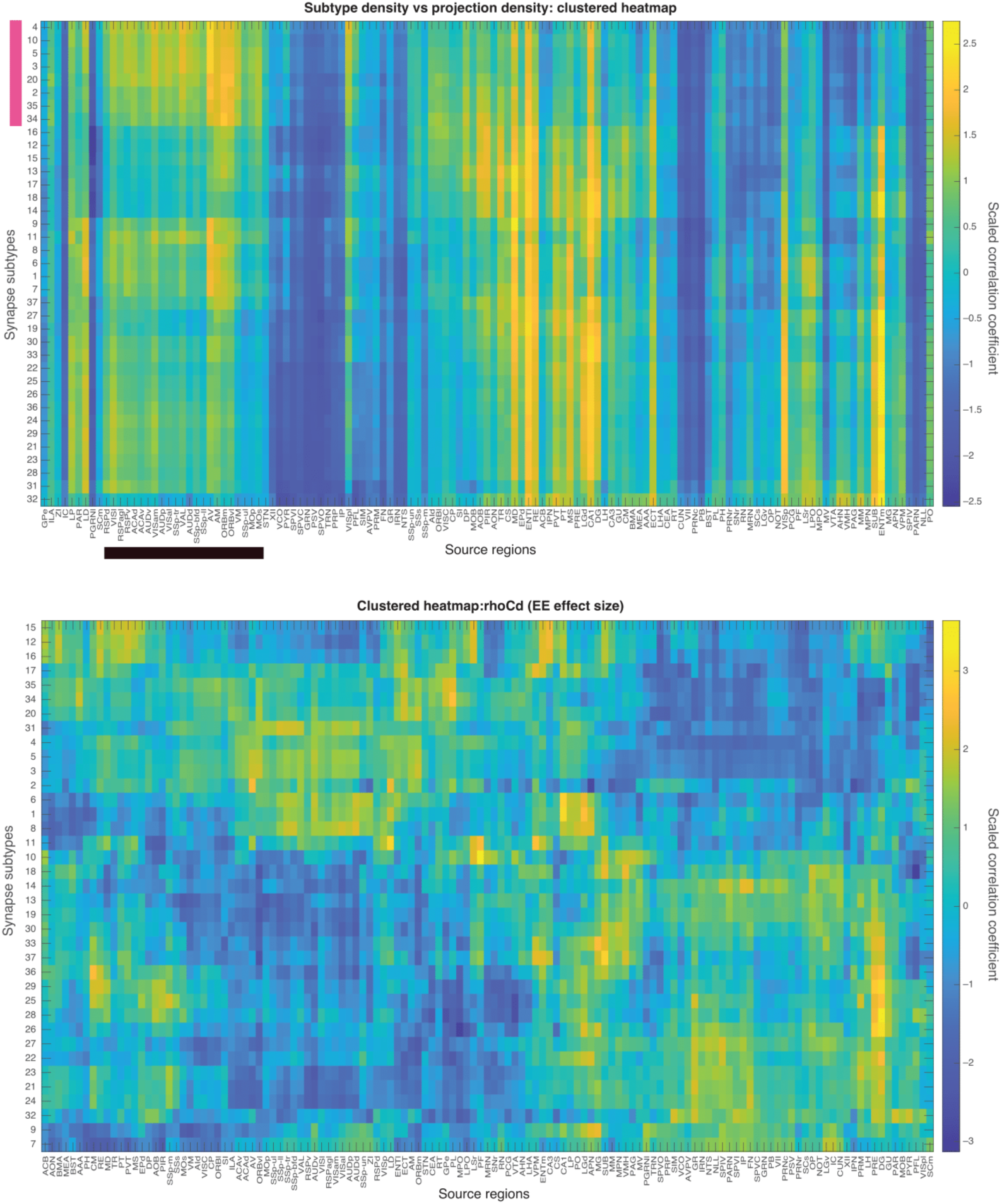
Top: Hierarchically clustered heatmaps of Spearman’s rho values correlating subtype densities (3-month control mice) with source region projection densities across target regions. The pink bar on the y-axis marks the cluster of long-protein lifetime synapses which positively correlate with several thalamocortical sensory areas, as indicated with the black bar on the x-axis. Bottom: Hierarchically clustered heatmaps of Spearman’s rho values correlating subtype effect size (developmental EE paradigm) with source region projection densities across target regions. Rows represent 37 synapse subtypes; columns are anatomically defined source regions. For brain region abbreviations see supplementary material ‘Brain region list’.

**Fig. S26).**
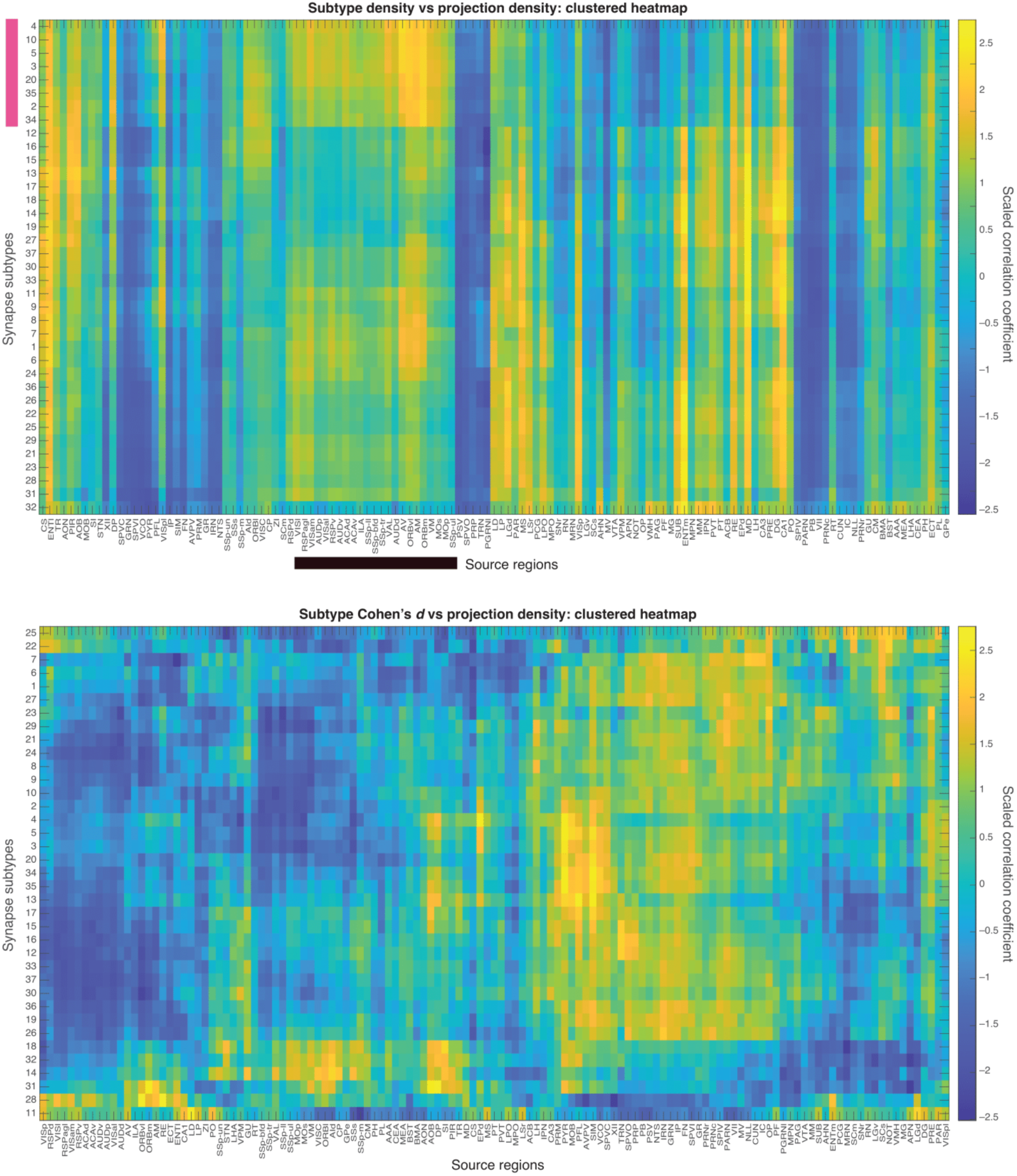
Top: Hierarchically clustered heatmaps of Spearman’s rho values correlating subtype densities (6-12-month control mice) with source region projection densities across target regions. The pink bar on the y-axis marks the cluster of long-protein lifetime synapses which positively correlate with several thalamocortical sensory areas, as indicated with the black bar on the x-axis. Bottom: Hierarchically clustered heatmaps of Spearman’s rho values correlating subtype effect size (adult EE paradigm) with source region projection densities across target regions. Rows represent 37 synapse subtypes; columns are anatomically defined source regions. For brain region abbreviations see supplementary material ‘Brain region list’.

**Fig. S27).**
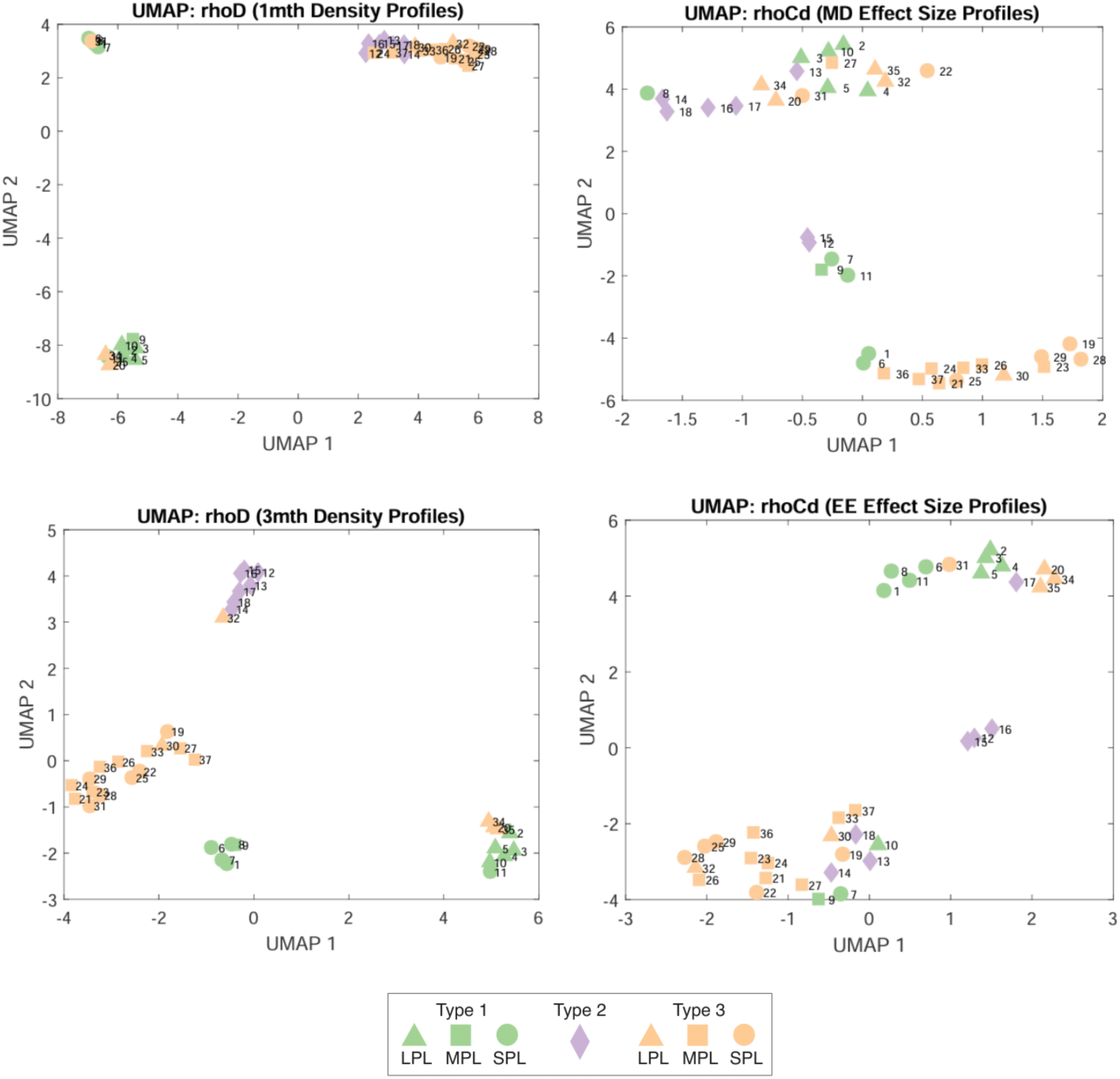
UMAP projections of synapse subtype correlation profiles with anatomical connectivity. UMAP was applied to z-scored Spearman’s rho correlation matrices linking projection density to either baseline subtype density (ρᴰ; top-left, bottom-left) or Cohen’s d effect size (ρᶜᵈ; top-right, bottom-right). (Top-left, bottom-left): UMAP projections of ρᴰ in 1-month (top) and 3-month (bottom) control mice. (Top-right, bottom-right): UMAP projections of ρᶜᵈ in MD (top) and developmental EE (bottom). Marker colour and shape indicate synapse type and protein lifetime class. UMAP parameters (k = 10 for MD, 8 for EE) were selected based on preservation of global structure (Mantel r: 0.66–0.79) and local fidelity (trustworthiness = 0.58–0.68; neighbour overlap = 0.62–0.71). Full embedding metrics for both UMAP and t-SNE are provided in the Supplementary Data.

**Fig. S28).**
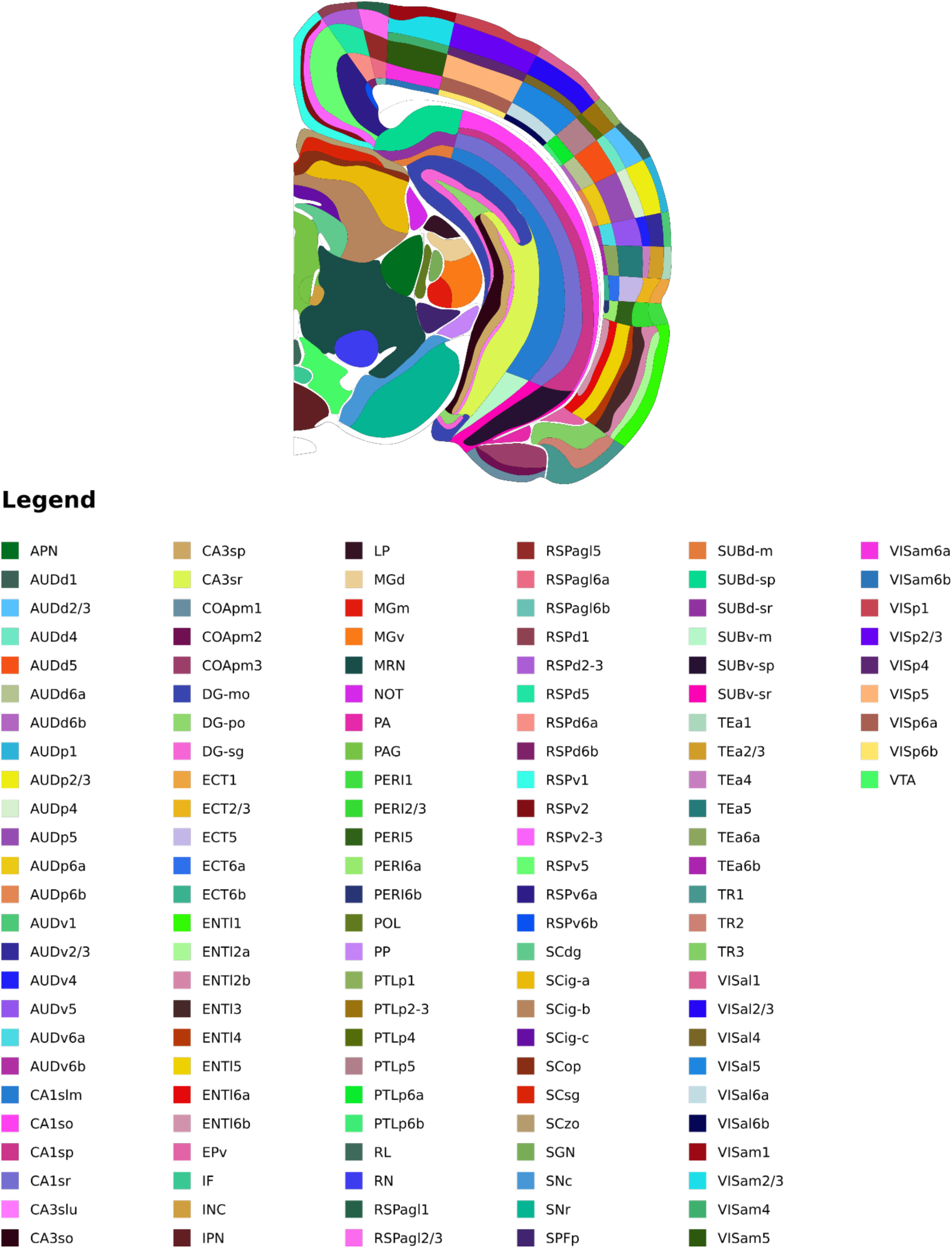
Coronal brain map coloured by subregion, as shown in legend. This figure was generated using a mapping tool developed in house based on brain atlases from the Allen Institute *(33)*. For brain region abbreviations see supplementary material ‘Brain region list’.

**Fig. S29).**
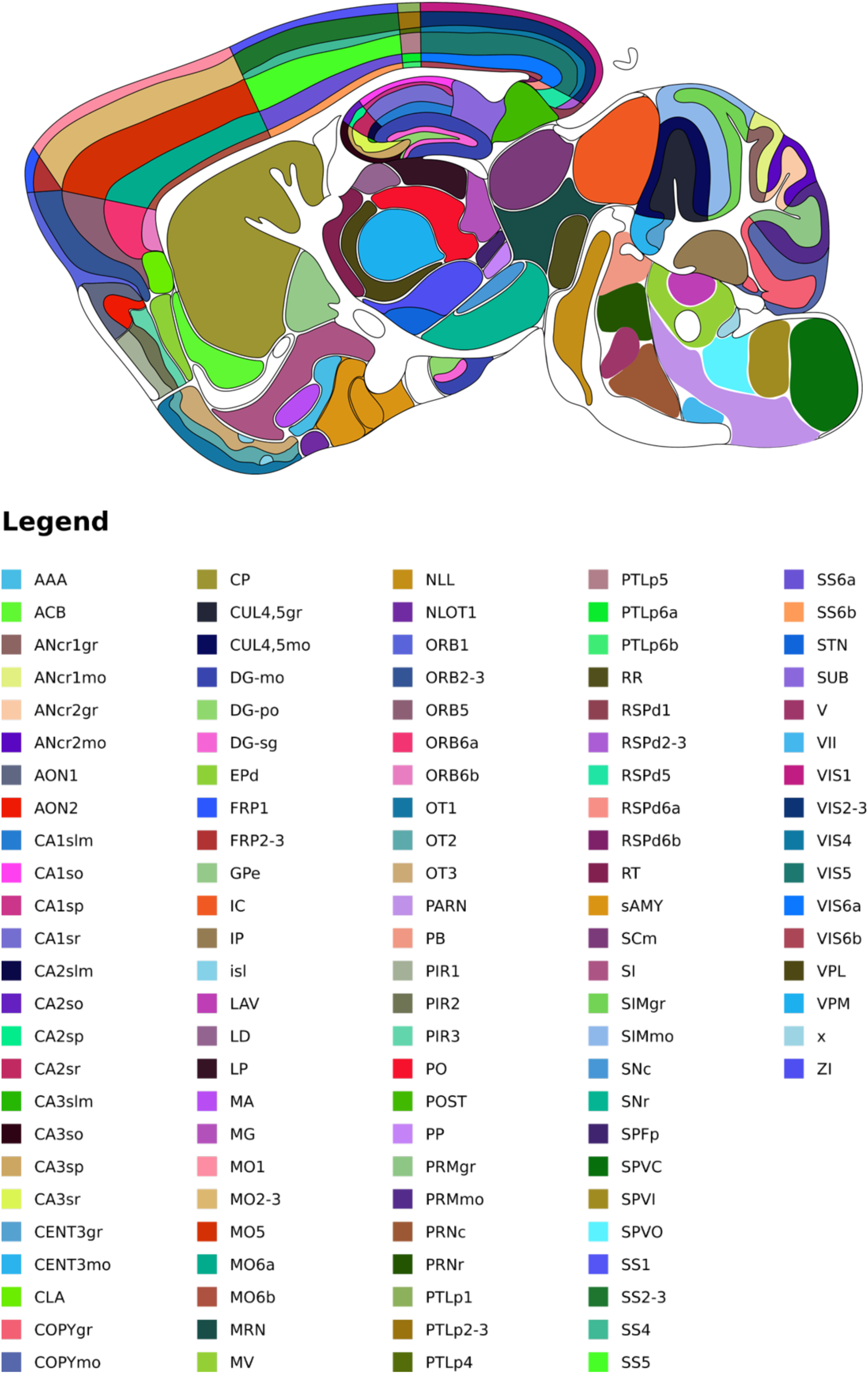
Parasagittal brain map coloured by subregion, as shown in legend. This figure was generated using a mapping tool developed in house based on brain atlases from the Allen Institute *(33)*. For brain region abbreviations see supplementary material ‘Brain region list’.

## Brain Region List

Acronym: Region name
ACB: Nucleus accumbens
AAA: Anterior amygdalar area
ACAd: Anterior cingulate area, dorsal part
ACAv: Anterior cingulate area, ventral part
ACB: Nucleus accumbens
AHN: Anterior hypothalamic nucleus
Aid: Agranular insular area, dorsal part
AM: Anteromedial nucleus of the thalamus
ANcr1gr: Crus 1, granular layer
ANcr1mo: Crus 1, molecular layer
ANcr2gr: Crus 2, granular layer
ANcr2mo: Crus 2, molecular layer
AOB: Accessory olfactory bulb
AON: Anterior olfactory nucleus
APN: Anterior pretectal nucleus
AUDd: Dorsal auditory area
AUDp: Primary auditory area
AUDv: Ventral auditory area
AV: Anteroventral nucleus of the thalamus
AVPV: Anteroventral periventricular nucleus
BMA: Basomedial amygdalar nucleus
BS: Brainstem
BST: Bed nuclei of the stria terminalis
CA1: Field CA1
CA1so: Field CA1, stratum oriens
CA1sp: Field CA1, stratum pyramidale
CA1sr: Field CA1, stratum radiatum
CA3: Field CA3
CA3slu: Field CA3, stratum lucidum
CA3so: Field CA3, stratum oriens
CA3sp: Field CA3, stratum pyramidale
CA3sr: Field CA3, stratum radiatum
CB: Cerebellum
CEA: Central amygdalar area
CENTgr: Lobule III, granular layer
CENTmo: Lobule III, molecular layer CLA Claustrum
CM: Central medial nucleus of the thalamus
COApm: Cortical amygdalar area, posterior part, medial zone
COPYgr: Copula pyramidis, granular zone
COPYmo: Copula pyramidis, molecular zone
CP: Caudate putamen
CS: Superior central nucleus raphe
CTX: Isocortex
CTXsp: Cortical subplate
CUL45gr: Lobules IV-V, granular layer
CUL45mo: Lobules IV-V, molecular layer
CUN: Cuneiform nucleus
DG: Dentate gyrus
DGmo: Dentate gyrus, molecular layer
DGpo: Dentate gyrus, polymorph layer
DGsg: Dentate gyrus, granule layer
DP: Dorsal peduncular area
ECT: Ectorhinal area
ENTl: Entorhinal area, lateral part
ENTm: Entorhinal area, medial part
EPd: Endopiriform nucleus, dorsal part
FN: Fastigial nucleus
FRP: Frontal pole
GPe: Global pallidus, external segment
GR: Gracile nucleus
GRN: Gigantocellular reticular nucleus
GU: Gustatory areas
HPF: Hippocampal formation
HY: Hypothalamus
IC: Inferior colliculus
ILA: Infralimbic area
IP: Interposed nucleus
IPN: Interpeduncular nucleus
IRN: Intermediate reticular nucleus
Isl: Islands of Cajella
LAT: Lateral group of the dorsal thalamus
LD: Lateral dorsal nucleus of the thalamus
LGd: Dorsal part of the lateral geniculate complex
LGv: Ventral part of the lateral geniculate complex
LH: Lateral habenula
LHA: Lateral hypothalamuc area
LP: Lateral posterior nucleus of the thalamus
LPO: Lateral preoptic area
LSr: Lateral septal nucleus, rostral part
LTD: Long-term depression
LTP: Long-term potentiation
MB: Midbrain
MD: Mediudorsal nucleus of the thalamus
MEA: Medial amygdalar area
MG: Medial geniculate complex
MM: Medial mammillary nucleus
MO: Somatomotor areas
MOB: Main olfactory bulb
Mop: Primary motor area
Mos: Secondary motor area
MPN: Medial preoptic nucleus
MPO: Medial preoptic area
MRN: Midbrain reticular nucleus
MS: Medial septal nucleus
MV: Medial vestibular nucleus
MY: Medulla
NLL: Nucleus of the lateral lemniscus
NOT: Nucleus of the optic tract
NTS: Nucleus of the solitary tract
OLF: Olfactory areas
OP: Olivary pretectal nucleus
ORB: Orbital cortex
ORBl: Orbital area, lateral part
ORBm: Orbital area, medial part
ORBvl: Orbital area, ventrolateral part
OT: Olfactory tubercle
P: Pons
PAG: Periaqueductal gray
PAL: Pallidum
PAR: Parasubiculum
PARN: Parvicellular reticular nucleus
PB: Parabrachial nucleus
PCG: Pontine central gray
PERI: Perirhinal area
PF: Parafascicular nucleus
PFL: Paraflocculus
PGRNl: Paragigantocellular reticular nucleus, lateral part
PH: Posterior hypothalamic nucleus
PIR: Piriform area
PL: Prelimbic area
PO: Posterior complex of the thalamus
POST: Post-subiculum
PP: Peripeduncular nucleus
PRE: Presubiculum
PRM: Paramedian lobule
PRMgr: Paramedian lobule, granular layer
PRMmo: Paramedian lobule, molecular layer
PRNc: Pontine reticular nucleus, caudal part
PRNr: Pontine reticular nucleus
PRP: Nucleus prepositus
PSV: Principal sensory nucleus of the trigeminal
PT: Parataenial nucleus
PTLp: Posterior parietal association areas
PVT: Paraventricular nucleus of the thalamus
PYR: Pyramus (VIII)
RE: Nucleus of reunions
RN: Red nucleus
RSPagl: Retrosplenial area, lateral agranular part
RSPd: Retrosplenial area, dorsal part
RSPv: Retrosplenial area, ventral part
RT: Reticular nucleus
RT: Reticular nucleus of the thalamus
sAMY: Striatum-like amygdalar nuclei
SCm: Superior colliculus, motor related
SCs: Superior colliculus, sensory related
SI: Substantia innominata
SIM: Simple lobule
SIMgr: Simple lobule, granular layer
SIMmo: Simple lobule, molecular layer
SNr: Substantia nigra, reticular part
SPFp: Subparafascicular nucleus, parvicellular part
SPIV: Spinal vestibular nucleus
SPVC: Spinal nucleus of the trigeminal, caudal part
SPVI: Spinal nucleus of the trigeminal, interpolar part
SPVO: Spinal nucleus of the trigeminal, oral part
SS: Somatosensory cortex
SSp-bfd: Primary somatosensory area, barrel field
SSp-ll: Primary somatosensory area, lower limb
SSp-m: Primary somatosensory area, mouth
SSp-tr: Primary somatosensory area, trunk
SSp-ul: Primary somatosensory area, upper limb
SSp-un: Primary somatosensory area, unassigned
SSs: Supplemental somatosensory area
STN: Subthalamic nucleus
SUB: Subiculum
SUBdm: Subiculum, dorsal part, molecular layer
SUBdsp: Subiculum, dorsal part, pyramidal layer
SUBdsr: Subiculum, dorsal part, stratum radiatum
SUBvm: Subiculum, ventral part, molecular layer
SUBvsp: Subiculum, ventral part, pyramidal layer
SUBvsr: Subiculum, ventral part, stratum radiatum
Tea: Temporal association areas
TH: Thalamus
TR: Postpiriform transition area
TRN: Tegmental reticular nucleus
VAL: Ventral anterior-lateral complex of the thalamus
VCO: Ventral cochlear nucleus
VII: Facial motor nucleus
VIS: Visual cortex
VISal: Anterolateral visual area
VISam: Anteromedial visual area
VISC: Visceral area
VISl: Lateral visual area
VISp: Primary visual area
VISpB: Primary visual cortex, binocular zone
VISpl: Posterolateral visual area
VISpM: Primary visual cortex, monocular zone
VM: Ventral medial nucleus of the thalamus
VMH: Ventromedial hypothalamic nucleus
VPL: Ventral posterolateral nucleus of the thalamus
VPM: Ventral posteromedial nucleus of the thalamus
VTA: Ventral tegmental area
XII: Hypoglossal nucleus
ZI: Zona Incerta

